# RBP45d is required for 5’splice site selection via binding to intronic U-rich elements and interaction with PRP39a in *Arabidopsis thaliana*

**DOI:** 10.1101/2022.08.12.503727

**Authors:** Weihua Huang, Liqun Zhang, Yajuan Zhu, Jingli Chen, Yawen Zhu, Fengru Lin, Jirong Huang

**Author notes:** Correspondence: Jirong Huang.

## Abstract

A large number of cryptic splice sites in eukaryotic genome are generally dormant unless activated by mutations of authentic splice sites or splicing factors. How cryptic splice sites are used remains unknown in plants. Here, we identified two cryptic splicing regulators, RBP45d and PRP39a that are homologs of yeast U1 auxiliary protein Nam8 and Prp39, respectively, via genetic screening for suppressors of the virescent *sot5* mutant, which results from a point mutation at the 5’ splice site (5’ ss) of intron 7. PCR and DNA sequencing data showed that loss-of-function mutations in *RBP45d* and *PRP39a* significantly increase the level of a cryptically spliced mRNA that encodes a mutated but partially functional sot5 protein, rescuing *sot5* to the WT phenotype. Yeast two hybrid and bimolecular fluorescence complementation assays demonstrated that RBP45d and PRP39a interact each other and also with the U1C, a core subunit of U1 small nuclear ribonucleoprotein (U1 snRNP). RNA electronic mobility shift assay showed that RBP45d directly binds to the uridine (U)-rich RNA sequence downstream of the cryptic 5’ ss. Consistently, our transcriptomic analysis revealed that a set of introns with U-rich sequences are retained in *rbp45d*. However, we found that other RBP45/47 members do not function redundantly with RBP45d, at least in regulation of cryptic splicing. Collectively, our data suggest that RBP45d is required for 5’ ss selection via binding to intronic U-rich elements and PRP39a in plants.

**One sentence summary:** The Arabidopsis RBP45d interacting with U1C and PRP39a is required for 5’ ss selection via binding to intronic U-rich elements.

## Introduction

Removal of introns from precursor mRNA (pre-mRNA), namely RNA splicing, plays an important role in regulation of gene expression at the posttranscriptional level in eukaryotes. It has been estimated that about 95% of human genes with multiple exons undergo alternative splicing and produce more than one mRNA variants, which significantly expands the human proteome (Chen and Manley, 2009). If pre-mRNA splicing is not accurately processed, it will lead to generation of various altered splicing events and subsequent nonfunctional proteins. In human genetic diseases, 10% are caused by abnormal splicing (Montes et al., 2019). Likewise, regulation of alternative splicing is also one of the important mechanisms regulating plant growth and development in responses to various environmental and developmental cues (Staiger and Brown, 2013; Reddyetal.,2013).

Pre-mRNA splicing is catalyzed by a macromolecular complex called spliceosome, which consists of five core small nuclear ribonucleoprotein particles (U snRNPs) and a number of auxiliary proteins (Chen and Manley, 2009). Coordination of U snRNPs with their auxiliary proteins and many non-snRNP proteins plays an important role in the recognition of splice sites and splicing efficiency. The intron usually contains a number of essential degenerate consensus motifs interacting with U snRNPs, such as at the 5’ and 3’ splice sites (ss), where the most conserved dinucleotides are GU and AG, respectively, and at the branch point site (BP), where A is present 20 to 50 nucleotides (nt) upstream of the 3’ss. In addition, exons and introns may contain *cis*-elements that can enhance or silence splicing and regulate constitutive splicing or alternative splicing via interaction with splicing regulatory factors (Chen and Manley, 2009, Lee and Rio, 2015). Interestingly, plant introns have evolved a number of distinct features from those in yeast and animals (Brown et al., 1996). For example, in plants, an intron is usually smaller in size and richer in A and U, the branch point sequence is not obvious, and the pyrimidine tract located between the BPS and 3’ss is mostly U (Brown et al., 1996). It has been demonstrated that the intronic AU-richness is essential for efficient splicing and 5’ ss selection in dicots (McCullough et al., 1993, Gniadkowski et al., 1996).

RNA splicing is initiated by base-pairing of the 5’ end of the U1 snRNA to the 5’ ss. In yeast and mammals, the 5’ ss consensus sequence for the major U2-type (GT-AG) introns can complement perfectly to that of the U1 snRNA (Roca and Krainer, 2008), which provides a mechanistic explanation for the important role of U1 snRNP in selection of 5’ ss. Besides U1 snRNA, U1 snRNP has seven core subunits called Sm proteins common in all the U snRNPs and three U1-specific proteins (U1-70K, U1A, and U1C) (Will and Luhrmann, 2011). It has been implicated that U1C is essential for U1 snRNP function by interacting with the Sm protein and recognizing the sequence of the 5’ ss (Nelissen et al., 1994; Du and Rosbash, 2002). In contrast, yeast *Saccharomyces cerevisiae* U1 snRNA (568 nts) is significantly larger than mammalian U1 snRNA (De Bortoli et al, 2019), and its U1 snRNP contains seven additional U1 auxiliary proteins including Prp39, Prp40, Snu71, Snu56, Luc7, Prp42, and Nam8 (Li et al., 2017). Among these U1 auxiliary proteins, the yeast Nam8 was reported to directly bind to the uridine (U)-rich sequence downstream of the 5’ ss and U1 snRNP at the same time, so as to effectively help splicing of introns with non-classical 5’ ss sequences (Puig et al., 1999). The human homologous protein of Nam8, TIA-1 (T-cell intracellular antigen-1) has been implicated in U1 snRNP recognition of the 5’ ss and alternative splicing regulation of various pre-mRNA via binding to U-rich motifs located downstream of 5’ ss and interaction with the U1C (Le Guiner et al., 2001; Forch et al., 2002). These data indicate that selection of 5’ ss is finely regulated not only by the complementary degree between the 5’ ends of the U1 snRNA and introns but also by many other factors, such as intronic U-rich elements and auxiliary splicing factors. Interestingly, there are eight Nam8 homologs, called the RNA binding protein (RBP) 45/47 family in *Arabidopsis* (Lorkovic et al., 2000; Peal et al., 2011). Recently, one member of the RBP45/47 family, RBP45d, was reported to associate with U1 snRNP via interacting with PRP39a and regulate alternative splicing (AS) for a specific set of genes (Chang et al., 2022). In addition, the Arabidopsis genome encodes many homologs of the seven yeast U1 auxiliary proteins (Wang and Brendel, 2004; Reddy et al., 2013). Thus, plant U1 snRNP has been suggested to be similar to or more complex than that in yeast. Although increasing genetic evidence supports that these U1 auxiliary proteins play an important role in plant growth, development and stress response (Wang et al, 2007; de Francisco Amorim et al., 2018; Hernando et al, 2019), molecular mechanisms by which they regulate U1 snRNP function remain largely unknown.

There are large numbers of splice sites, known as cryptic splice sites in Eukaryotic genomes, which are generally dormant or used only at low levels unless activated by mutations of nearby authentic splice sites (Roca et al., 2003; Kapustin et al, 2011). Notably, about 9% of all mutations reported in the human gene mutation database are splicing mutations (Annal and Monika 2018). It is important to be able to predict cryptic splice sites that might be activated in genetic disease so that effective therapeutic strategies can be designed. On the other hand, mutations leading to activation of cryptic splice sites in crops have generated new genetic germplasm resources, such as *waxy* rice, low phytic acid soybean, and pale green Chinese cabbage (Isshiki et al., 1998; Yuan et al., 2012; Zhao et al., 2022). It is generally accepted that cryptic splice sites are suppressed by nearby stronger splice sites and that splice site selection can be viewed as a competition between the various potential splice sites in a pre-mRNA (Kapustin et al, 2011). And it is also proposed that cryptic splice sites are regulated by various RNA binding proteins including heterogeneous ribonucleoproteins (hnRNP) and RNA recognition motif (RRM)-containing SR (serine and arginine-rich) proteins, which are also the alternative splicing regulators (Annal and Monika 2018). Little is known about splicing factors involved in regulating cryptic splicing in plants. In this study, we showed that Arabidopsis RBP45d facilitates U1 snRNP to preferentially select a cryptic splice site that is activated by mutations of the intronic 5’ ss via directly binding to U-rich sequences downstream of the 5’ ss. Our results suggest that RBP45d regulates alternative splicing and cryptic splice site selection in plants.

## Results

### An intron mutation leads to activation of two cryptic splice sites in *sot5*

We previously reported that the *Arabidopsis* PPR protein SOT5 (At1g30610), also named EMB2279, is required for intron splicing of the two chloroplast housekeeping genes, *rpl2* and *trnk* (Huang et al., 2018). The *sot5* or *emb2279-3* mutant contains a G to A mutation at the first base (+1) of the seventh intron and displays a leaf virescent phenotype (Figure 1A and 1C). This point mutation leads to accumulation of the three alternative splicing variants in *sot5*, compared to the wild type (WT) (Figure 1B). DNA sequencing of these cloned PCR products indicated that the longest one retained the whole seventh intron with 85 nts (Figure 1B, band b and Figure S1), the middle transcript retained 9 nts more at the 3’ ss of exon 6 (Figure 1B, band c and Figure S1), and the shortest one lacked 22 nts at the 3’ end of exon 6 (Figure 1B, band d and Figure S1). These results indicate that the mutation of the first invariant G at the 5’ ss of intron 7 results in intron retention and activation of two cryptic splice sites named −22 ss and +9 ss (Figure 1C). Since the level of band d was much higher than that of band c, the cryptic splicing mainly occurred at −22 ss in *sot5* (Figure 1B). Polypeptide prediction indicated that the −22 ss product encodes a truncated protein with only 6 PPR motifs that should be loss of function (Figure 1D); the intron-retained transcript encodes a probably partially functional protein with 10 PPR motifs (Chen et al, 2020); the +9 ss product encodes almost the same protein as the endogenous one except the seventh PPR motif is slightly changed (Figure 1D, Figure S2). Thus, these results interpret well why *sot5* displays a virescent phenotype, but the *emb2279-1* knockout mutant is embryo lethal.

**Figure 1.**
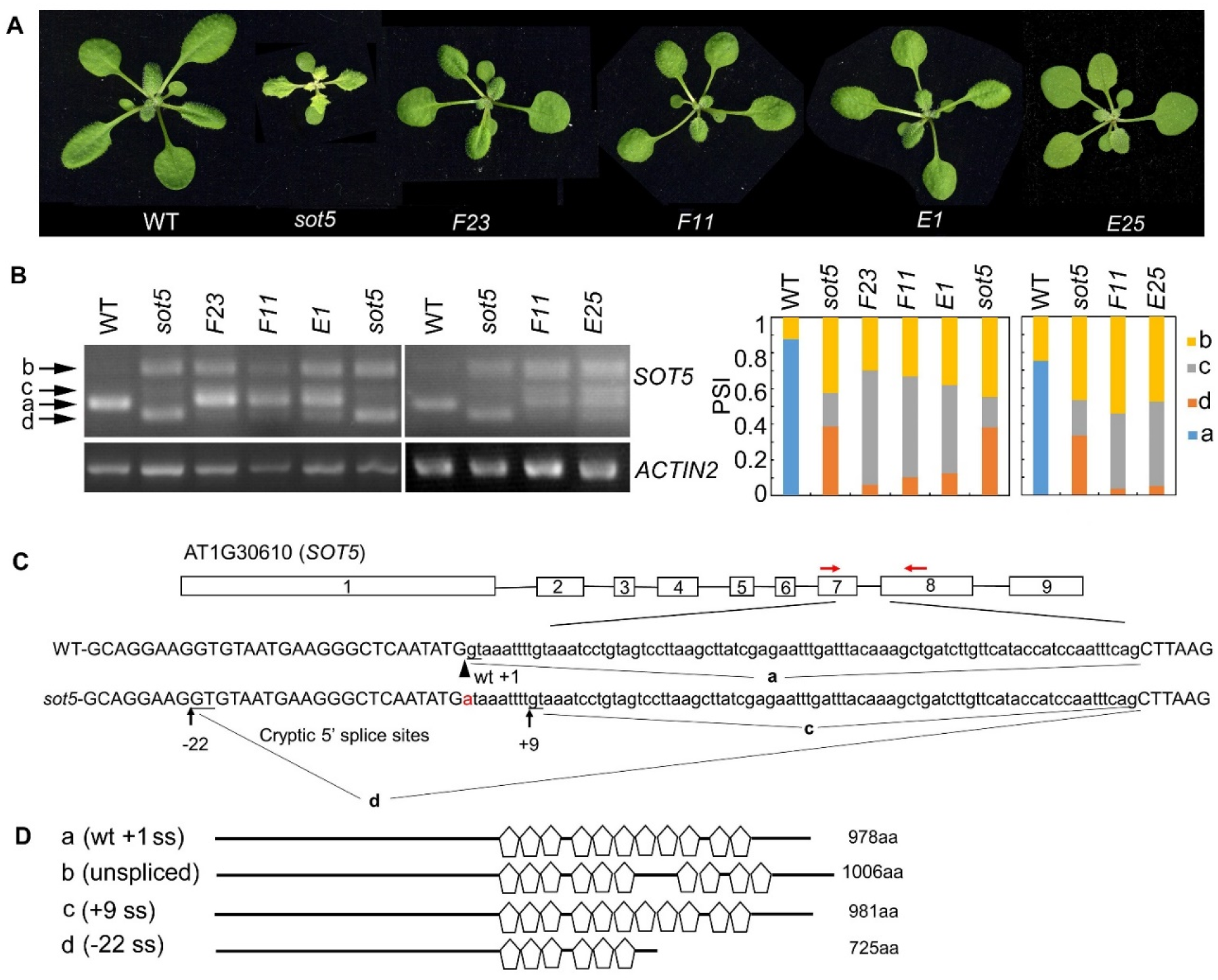
Genetic screening for suppressors of the *sot5* viresent phenotype reveals altered cryptic 5’ splice sites by second mutations. (A) Four suppressor lines of *sot5* display WT-like phenotype. The representative 30-d-old plants were shown. (B) The left panel shows the splicing products of *SOT5* intron 7 in WT, *sot5*, and the suppressor lines. *ACTIN2* was used as an internal control. Band a shows the WT splicing product. Band b, c, and d correspond to the splicing variants containing intron 7, derived from splice site +9 ss and −22 ss, respectively. The right panel shows the quantification of the four bands in each genotype using Image J software. PSI (percent spliced in index) indicates the percentage of each variant in total transcripts. Three biological replicates (independent pools of aerial parts of plants) were analyzed, and one representative result is shown. (C) The schematic diagram of *SOT5* gene structure and intron 7 splicing variants. The black boxes indicate exons and the black lines indicate introns. The authentic 5’ ss in WT is shown by the arrowhead. The G to A point mutation in *sot5* is shown by the red letter. Black arrows indicate two cryptic splice sites, +9 ss and −22 ss. a, c and d indicated the splicing variants corresponding to the bands in (B). Red arrows indicate the primers used for RT-PCR. (D) Schematic diagrams of the mutated SOT5 proteins encoded by the variants corresponding to the bands in (B). Pentagons indicate PPR motifs. The numbers indicate the residue number of each protein.

### Genetic screening for suppressors of *sot5* reveals altered cryptic splicing by second mutations

The weak allele *sot5*, in which an intron mutation leads to generation of three abnormal transcripts, provides a good material to study regulation of the cryptic splice site. To dissect the genetic pathway for the cryptic splicing of *SOT5* intron 7, we screened for *sot5* suppressors which exhibit a WT-like phenotype using the ethyl methanesulfonate (EMS)-mutagenized *sot5* seeds. We obtained a number of WT-like suppressor lines for the *sot5* mutant (Figure S3). In this study, four suppressor lines, namely *F23*, *F11*, *E1* and *E25*, were selected for further analysis based on their similar phenotype and alternative splicing pattern of *SOT5* intron7 (Figure 1A, 1B and Figure S3). Compared to *sot5*, all these suppressor lines produced a lower level of −22 ss transcripts (band d in Figure 1B), but a higher level of +9 ss transcripts (band c in Figure 1B), suggesting a positive correlation between the green leaf and the level of the +9 ss transcripts. In addition, our genetic analyses showed that the suppressor phenotype was caused by a single gene mutation, and interestingly *F23, F11* and *E1* were allelic while *E25* was mutated in another gene. Taken together, our data suggest that the *sot5* phenotype is suppressed by second mutations that can modulate cryptic splice site selection and splicing efficiency.

#### Loss-of-function mutations in *RBP45d* suppress the *sot5* phenotype

To isolate the suppressor gene in *F23*, we produced a mapping population from the cross between *F23* and Landsberg *erecta* (L*er*). Through Mapping-By-Sequencing (MBS) technique, the gene responsible for phenotypic suppression of *sot5* was mapped to the region with the length ~8 M bp on chromosome 5, which contains 7 point mutations (Figure S4A and S4B). Among these 7 mutations, one stop codon-gain mutation from C to T occurred at 187 bp of the CDS in the locus At5g19350, which encodes RBP45d, a member of the RBP45/47 family with triple RNA recognition motifs (RRM) (Figure 2A and S1B). Considering that *sot5*-activated cryptic splicing efficiency is altered in *F23*, we speculated that *RBP45d* was the most likely gene to suppress *sot5*. Thus, we sequenced the *RBP45d* gene in *F11* and *E1*, which were genetically testified as *F23* alleles. Indeed, *F11* has a point mutation from G to A at the 481 bp position from the start codon of the *RBP45d* CDS, resulting in the substitution of glycine (G) by arginine (R) at the 161 amino acid residue in the conserved RRM2 domain, whereas *E1* contains a premature stop codon mutation from G to A at the 80 bp position from the start codon of the *RBP45d* CDS (Figure 2A).

**Figure 2.**
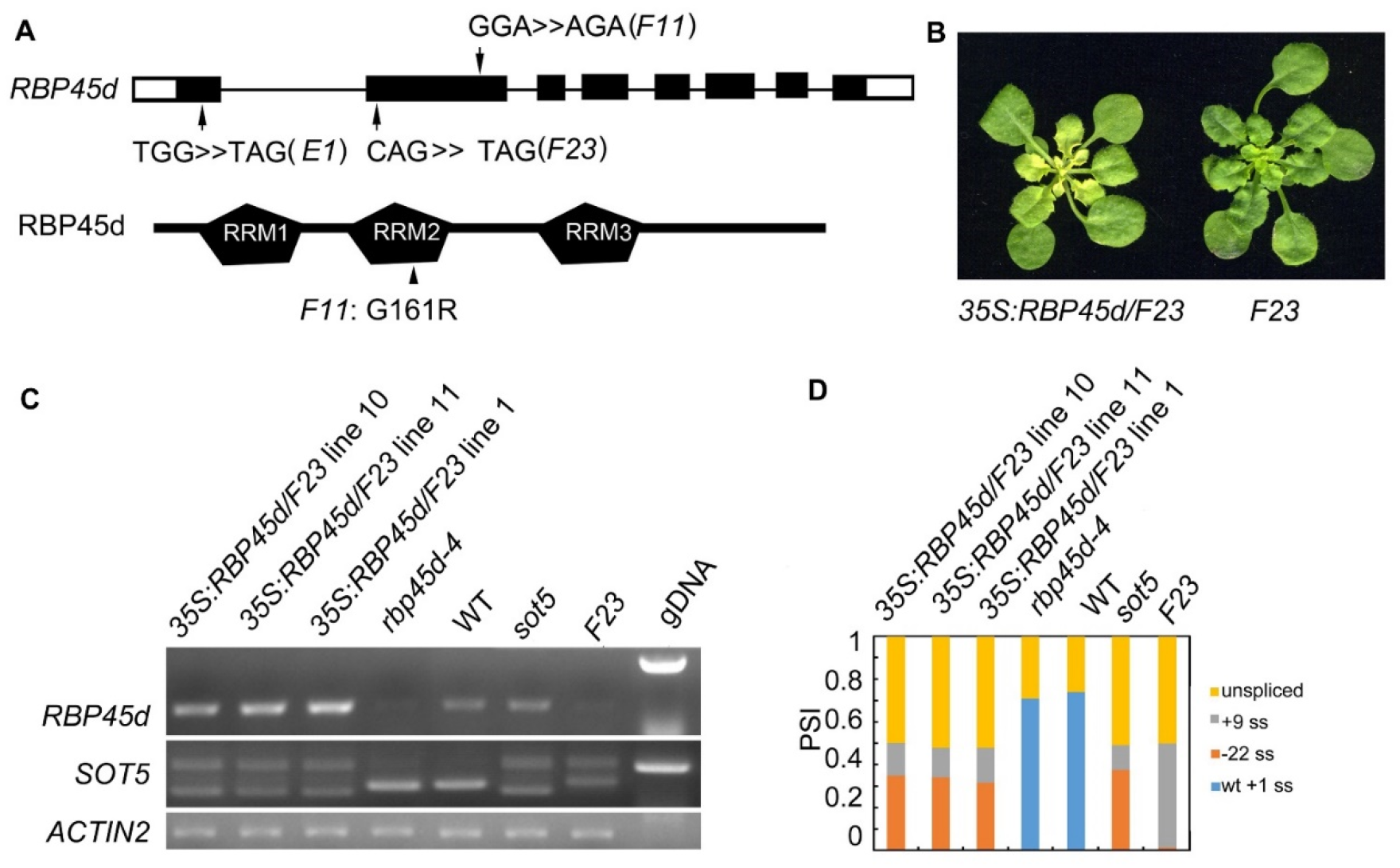
Mutations in RBP45d suppress the *sot5* phenotype. (A) Schematic diagrams of *RBP45d* gene structure and RBP45d protein with the three RRM domains. Arrows indicate the mutation sites in *E1*, *F23* and *F11* lines. The upper case letters indicate the mutated genetic codon(s) in *E1* and *F23* which become the stop codons, and that in *F11* which leads to replacement of the amino acid residue from G to R in the conservative region of the RRM2 domain (arrowhead). The black boxes indicate exons in UTR and the black rectangles indicate exons in CDS and the black lines indicate introns. (B) Overexpression of *RBP45d* CDS in *F23* restores the phenotype. One of representative *35S:RBP45d/F23* line was shown. (C) RT-PCR analysis of *RBP45d* expression and the splicing products of *SOT5* intron 7 in three *35S:RBP45d/F23* transgenic lines, *rbp45d-4*, WT, *sot5*, and *F23* plants. *ACTIN2* was used as an internal control and gDNA is used as a control. Two biological replicates (independent pools of aerial parts of plants) were analyzed, and one representative result is shown. (D) Quantification of the splicing variants from *SOT5* intron 7 shown in (C). PSI (percent spliced in index) indicates the percentage of each variant in total transcripts.

To verify the mapping result, we overexpressed the native *RBP45d* CDS in *F23*. Our data showed that transgenic lines *35S:RBP45d*/*F23* not only displayed the *sot5* phenotype (Figure 2B), but also had the same alternative splicing pattern as the *sot5* mutant (Figure 2C and 2D), indicating that the phenotypic recovery of *sot5* is attributed to the loss-of-function mutation in *RBP45d*. We then isolated the single mutant, named *rbp45d-4*, from the F_2_ population of the cross between *F23* and Col-0 and *rbp45d-5* from the F_2_ population of the cross between *F11* and Col-0. RT-PCR analysis showed that *RBP45d* transcripts were almost undetectable in *rbp45d-4* and *F23* (Figure 2C), indicating that the point mutation in *rbp45d-4* triggers the nonsense-mediated mRNA decay (NMD) pathway. Taken together, our results imply that *RBP45d* mutations are responsible for the recovery of *sot5* leaf virescence.

#### Mutations of other *RBP45/47* members cannot suppress the phenotype of *sot5*

It has been reported that the *Arabidopsis* genome has eight members in the RBP45/47 family (Wang and Brendel, 2004). RT-PCR analysis and eFP Browser (http://bar.utoronto.ca/efp_arabidopsis/cgi-bin/efpWeb.cgi) showed that they are ubiquitously expressed in all organs including roots, inflorescences and leaves (Figure S5). Subcellular localization experiment showed that RBP45a, RBP45b, RBP45c, RBP45d and RBP47b were localized in the both nucleus and cytoplasm, while RBP47c’ mainly localized in the cytoplasm (Figure S6). The mutant phenotype and physiological function of RBP45/47 family members were not well characterized yet. To know whether other members of the RBP45/47 family can function in the same manner as RBP45d to suppress the *sot5* phenotype, we identified four T-DNA insertion mutants *rbp45a, rbp45c, rbp47a* and *rbp47b* from the ABRC stock (Figure 3A and 3B), and made their double mutants with *sot5* by genetic crossing. All the four single mutants displayed the WT-like phenotype under normal growth conditions, while the double mutants with *sot5* showed the *sot5*-like phenotype (Figure 3C). Therefore, our data indicate that *rbp45a, rbp45c, rbp47a* and *rbp47b* cannot suppress the phenotype of *sot5*.

**Figure 3.**
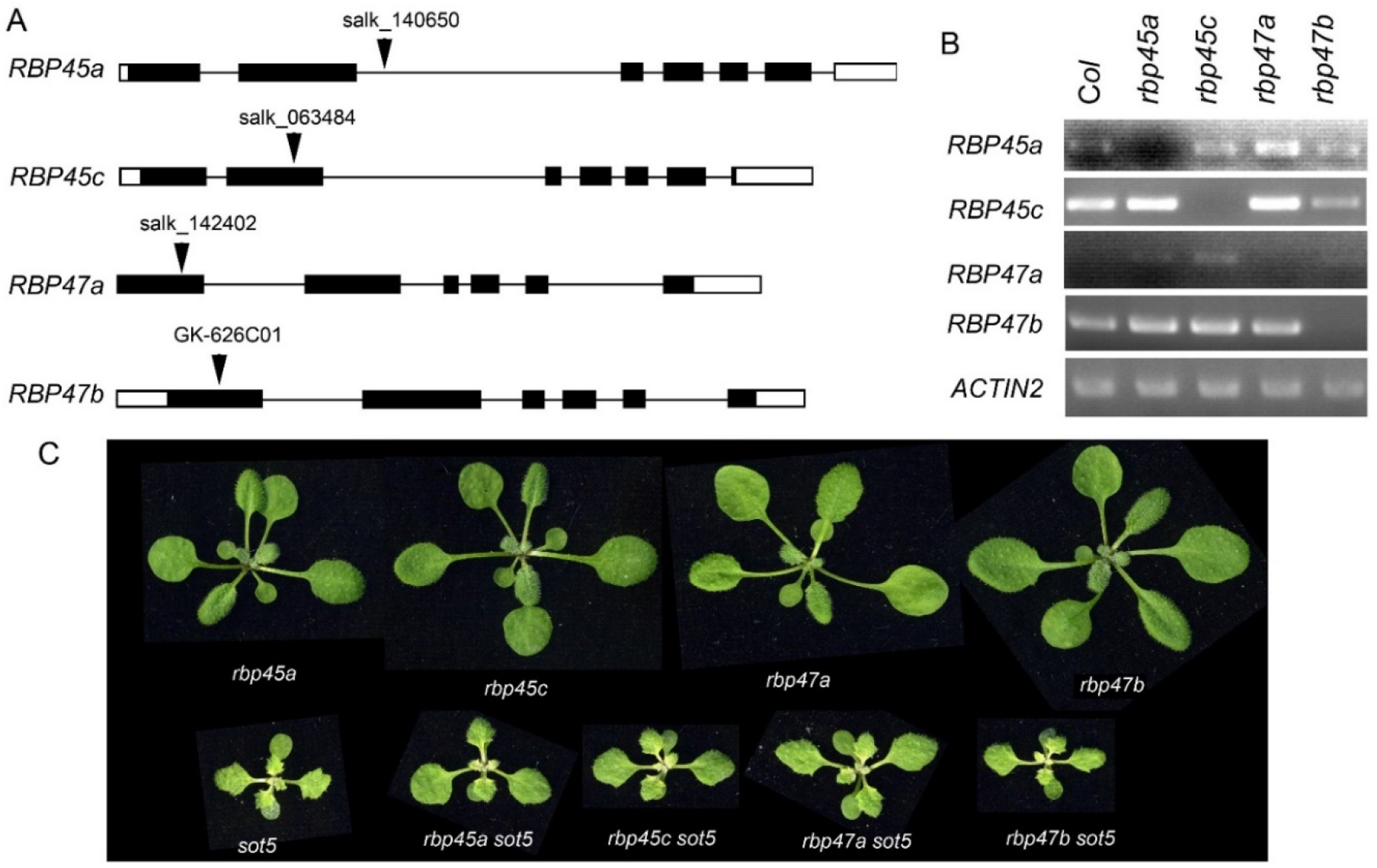
Mutations in other *RBP45/47* paralogs cannot suppress the viresent phenotype of *sot5*. (A) Identification of T-DNA insertions in *RBP45a, RBP45c, RBP47a* and *RBP47b* genes. The black boxes indicate exons in UTR and the black rectangles indicate exons in CDS and the black lines indicate introns. Arrowheads indicate the insertional position of T-DNA for each gene. (B) RT-PCR analysis of gene expression in the T-DNA knockout mutants. *ACTIN2* was used as an internal control. Two biological replicates (independent pools of aerial parts of plants) were analyzed, and one representative result is shown. (C) Phenotypes of single and double mutants as indicated.

### RBP45d functions differentially from other *RBP45/47* members in plant growth and development

Previously, Chang et al (2022) reported that *RBP45d* regulates temperature-responsive flowering in *Arabidopsis*. In this study, we observed the same later flowering phenotype of *rbp45d* mutants including *rbp45d-4, rbp45d-5* and *rbp45d-CR* (a knockout *rbp45d* mutant created by CRISPR/CAS9 technique) under our growth conditions (Figure 4A upper panel and Figure S7). The number of rosette leaves for *rbp45d* mutants to flower were significantly higher than that for WT (Figure 4B the left panel and Figure S7B). However, there were no significant difference in flowering time between WT and other *rbp45/47* plants (Figure 4A and 4B). In addition, we found that loss-of-function mutations in *RBP45d* but not in other *RBP45/47* genes led to the shorter primary root phenotype, compared to WT, when seedlings were vertically grown on 1/2 MS plates (Figure 4A lower panel, Figure S7). For 5-day-old seedlings, primary root length of *rbp45d-4* and *rbp45d-5* was 64% and 80% of WT, respectively (Figure 4A lower panel). Such a difference in primary root length became much larger between 10-day-old *rbp45d* and WT or other *rbp45/47* seedlings. These results suggest that the function of RBP45d is different from that of other RBP45/47 paralogs.

**Figure 4.**
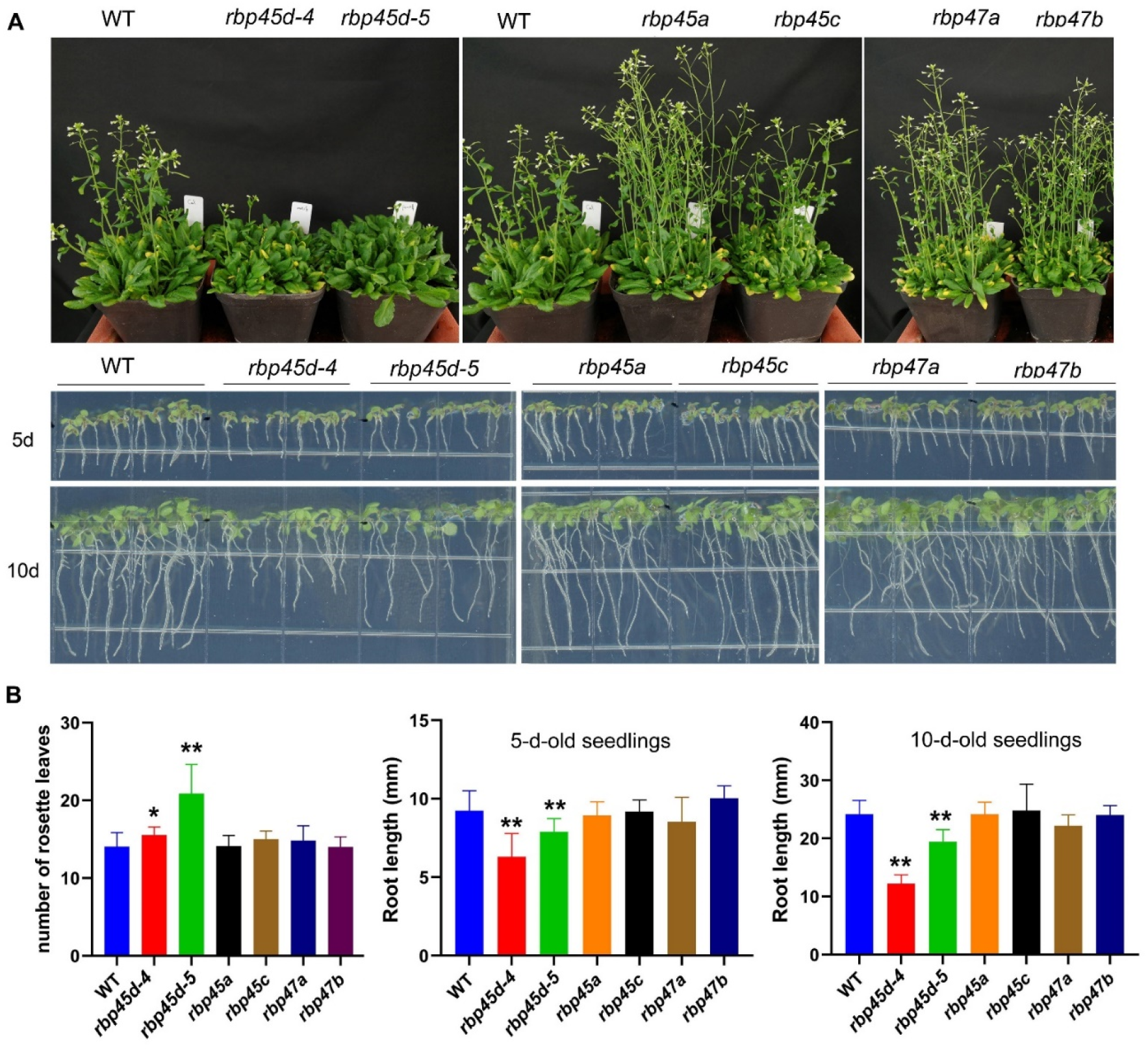
Phenotype of *rbp45d* mutant is distinct from that of other *rbp45/47* mutants. (A) Phenotypic comparison among *rbp45/47* mutants. The upper panel shows the phenotype of 45d-old *rbp45/47* mutants grown under LD conditions (16h light/8h dark photoperiod). The lower panel shows 5-d- and 10-d-old *rbp45/47* mutants vertically grown on 1/2 MS plates. (B) The left panel shows the rosette leaf number of each genotype at bolting in LD conditions. The middle and right panels show root length of 5-d and 10-d-old seedlings, respectively. The data are shown means ± SD (n = 20). Student’s *t*-test, * *P* < 0.05, ** *P* < 0.01, compared to WT. The experiments were performed at least three times.

### RBP45d physically interacts with the U1 snRNP core subunit U1C

RBP45/47 proteins are homologous with yeast Nam8p protein and human TIA-1 protein. It was reported that TIA-1 can directly interact with U1C, the core protein of U1 snRNP (Forch et al., 2002). And the cryo-electron microscopy structure of the yeast *Saccharomyces cerevisiae* prespliceosome revealed that yeast Nam8 protein physically interacts with the U1C. Thus, we proposed that RBP45d may function in a similar manner with TIA-1 and Nam8. We then used yeast two hybrid (Y2H) and BiFC techniques to detect whether RBP45d interacts directly with U1C, U1 70k and U1A of U1 snRNP in *Arabidopsis*. Y2H results showed that RBP45d directly interacted with U1C, but not with U1A and U1 70K (Figure 5A). Consistently, BiFC analysis also revealed that RBP45d was bound to U1C in both the nucleus and cytoplasm in *Arabidopsis* protoplasts (Figure 5B), which is consistent with their subcellular localization (Figure S6). Taken together, these results indicate that RBP45d interacts with U1C *in vitro* and *in vivo*.

**Figure 5.**
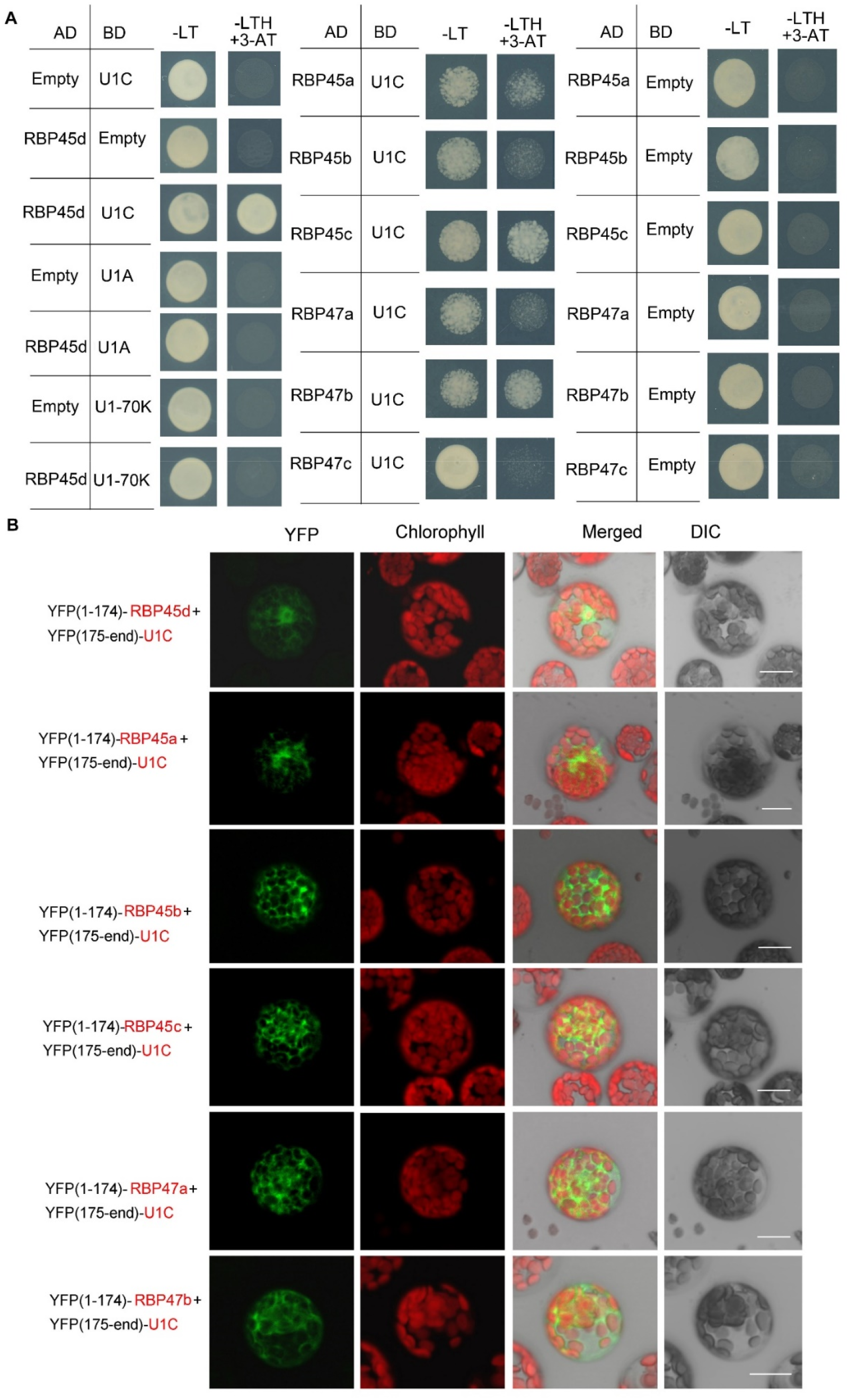
RBP45d and other RBP45/47 members interact with U1C. (A) Yeast two-hybrid assays show the interaction between RBP45d and U1C, U1A or U1 70K and the interaction between U1C and RBP45a, RBP45b, RBP45c, RBP47a, RBP47b or RBP47c. The transformed yeast cells were grown on −LT (SD-Leu/-Trp) mediums and −LTH+3-AT (SD-Leu/ −Trp /-His with 5 mM 3-AT) mediums. (B) BiFC assay of the interaction between U1C and RBP45d, RBP45a, RBP45b, RBP45c, RBP47a or RBP47b. YFP, fluorescence of yellow fluorescent protein; DIC, Differential interference contrast microscopy; Merge, merge of YFP, DIC and chlorophyll. Bar = 15 μm.

Meanwhile, Y2H analysis showed that except for RBP47C, the RBP45/47 members interacted with U1C (Figure 5A), whereas BiFC assays indicated that all the tested RBP45/47 paralogs interacted with U1C mainly in the cytoplasm (Figure 5). Thus, our results indicate that the biological significance of U1C interaction with RBP45d may differs from that with other RBP45/47 members.

### RBP45d is an intronic U-rich element binding factor

The above results suggest that RBP45d is recruited to the U1 snRNP via interacting with U1C, and prompted us to address whether RBP45d can interact with U-rich sequences. In the *sot5* mutant, the G to A mutation at the 5’ ss of intron 7 results in activation of the two cryptic splice sites (−22 ss and +9 ss) and preferential utilization of the −22 ss (Figure 1B and 1C). In contrast, the absence of RBP45d in the suppressor lines *F23*, *F11* and *E1* leads to preferentially utilize +9 ss (Figure 1B), implying that RBP45d plays an important role in controlling alternative splicing efficiency between the cryptic splice sites. Search for U-rich elements within *sot5* intron 7 indeed revealed the presence of the AUAAAUUUU sequence downstream of the −22 ss (Figure 6A). We proposed that RBP45d can bind to this U-rich sequence and subsequently promote the upstream −22 ss selection.

**Figure 6.**
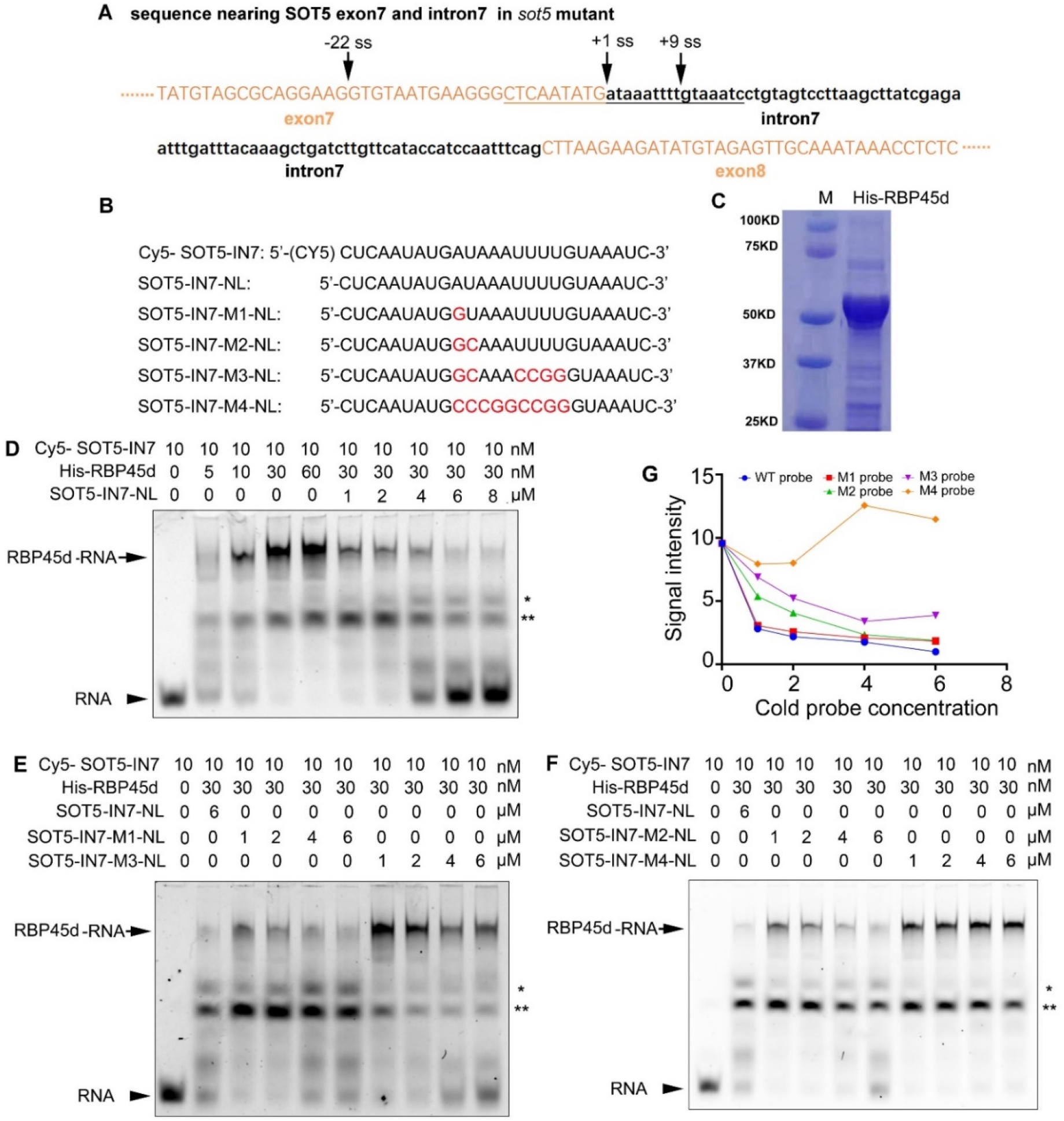
REMSA shows direct binding of RBP45d to the U-rich sequence downstream of the −22 ss of *SOT5* intron 7. (A) DNA sequence nearby the cryptic 5’ splice sites in *sot5*. Upper and lower case letters indicate exonic and intronic sequences in *SOT5* gene, respectively. The underlined sequence is used as a template for RNA probe synthesis. Arrows, splice sites. (B) Probe names and sequences used for REMSA. The mutation sites in the probes were highlighted with red color. (C) Coomassie brilliant-blue staining of the purified His-RBP45d protein separated by SDS-PAGE. (D) RBP45d protein can directly bind to the Cy5-SOT5-IN7 RNA probe. (E) Competitive binding assay of unlabeled probes SOT5-IN7-M1-NL and SOT5-IN7-M3-NL with the labeled Cy5-SOT5-IN7 (WT) probe. (F) Competitive binding assay of unlabeled probes SOT5-IN7-M2-NL and SOT5-IN7-M4-NL with the labeled WT probe. Arrows in (D), (E), and (F) show the protein-RNA complex. Arrowheads in (D), (E), and (F) show the free RNA probe. * and ** in (D), (E), and (F) indicate non-specific bands. (G) Relative signal intensity of the protein-RNA complex in (D), (E), and (F) was quantified by ImageJ software. The signal intensity of the protein-RNA band in the lane using 6 μM unlabeled SOT5-IN7-NL to compete with the labeled Cy5-SOT5-IN7 probe is artificially set to 1 in (D), (E), and (F). For REMSA assays, three technical replicates were analyzed, and the results have the same trend. So one representative result is shown.

To test this hypothesis, we carried out RNA electrophoretic mobility shift assay (REMSA) with a series of synthesized 26 nt RNA oligo probes including the WT U-rich element (SOT5-IN7) and mutated variants (SOT5-IN7-Ms), and purified His-tagged RBP45d protein from *E. coli* (Figure 6B and 6C). REMSA results showed that RBP45d bound to the WT probe (Cy5-SOT5-IN7) in a dose-dependent manner, and the unlabeled WT probe (SOT5-IN7-NL) was able to compete with the labeled probe in RBP45d binding (Figure 6D). Further analysis showed the unlabeled SOT5-IN7-M1 and SOT-IN7-M2 probes, in which the first A and AU of intron 7 were mutated into G and GC, respectively, still efficiently blocked RBP45d binding to the labeled WT probe (Figure 6E and 6F), whereas the unlabeled SOT5-IN7-M3 probe with mutations at the first AU and 4 continuous U of intron 7 significantly reduced the binding affinity to RBP45d (Figure 6E). In contrast, the SOT5-IN7-M4 probe completely lost the binding affinity to RBP45d when the AU-rich element was mutated into the CG-rich one (Figure 6F). Quantification of the signal intensity of these protein-RNA bands revealed that the affinity of RBP45d binding to RNA increased with the increase in U and/or A levels of the probes (Figure 6G). Thus, our results suggest that RBP45d can bind to the U-rich sequence element, and binding affinity between RBP45d and RNA is dependent on the content of U and/or A.

### *RBP45d* mutations lead to retention of the introns with the U-rich element

To identify the RBP45d-biding *cis* element *in vivo*, we first performed high-throughput transcriptomic assay with the two pairs of samples, namely *rbp45d* and Col, and *F23* and *sot5*. Our results showed that *RBP45d* mutations led to a significant effect on intron retention, despite affecting all types of splicing events, including alternative 5’ ss, alternative 3’ ss, alternative start exon, alternative end exon, and mutually exclusive exons (Figure 7A and 7B, Table S1). A total of 109 and 365 alternative splicing events were significantly altered in *rbp45d* and *F23* (FDR < 0.05), compared to their corresponding control, WT and *sot5*, respectively (Table S2 and S3). Venn diagram analysis (http://www.ehbio.com/test/venn/) showed that 50 alternative splicing events were overlapped in the two groups (Figure 7C, Table S4 and S5). Among these 50 alternative splicing events, 23 intron retention events significantly increased in both *rbp45d* and *F23* (delta_PSI > 0) (Table S5 and S6). We then verified some of the 23 intron retention events in *rbp45d* and WT plants by RT-PCR (Figure 7D). As shown in Figure 7D, compared to those in the WT, the examined introns transcribed from *mitochondrial lipoamide dehydrogenase1* (*mtLPD1*), *Scream2* (*SCRM2*), *Uroporphyrinogen decarboxylase* (*HEME1*), Sinapoylglucose1 (*SNG1*), *Beta glucosidase8* (*BGLU8*), Cysteine synthase C1 (*CYSC1*), *at3g51950*, and *at2g45990* were significantly retained in *rbp45d-4*. Subsequently, we searched for a consensus sequence element in these 23 introns by the MEME suite (https://meme-suite.org), and found the conserved U-rich element (Figure 7E, Table S6). Lastly, we investigated whether RBP45d protein can directly bind to the U-rich sequence. To do this, we took the second intron of *mtLPD1* as an example, and synthesized the 29 nt RNA probe containing the U-rich elements, named LPD1-IN2 (Figure 7F). REMSA results showed that LPD1-IN2 binding affinity to RBP45d increased with an increase in the amount of RBP45d protein, indicating that RBP45d protein strongly bind to LPD1-IN2 RNA probe. We also evaluated different RBP45d-binding affinity between LPD1-IN2 and SOT-IN7 probes. Interestingly, the unlabeled SOT5-IN7 probe could not completely suppress RBP45d binding to the labeled LPD1-IN2 probe although the concentration of probe SOT5-IN7 was 150 times of the LPD1-IN2 concentration (Figure 7F), indicating that the affinity of RBP45d to *LPD1* intron 2 probe is much higher than that of SOT5-IN7 probe. This probably caused by the higher U content of LPD1-IN2 probe than SOT5-IN7 probe. Taken together, our data suggest that RBP45d protein can directly bind to the intronic U-rich element downstream of the 5’ ss and enhance intron splicing efficiency.

**Figure 7.**
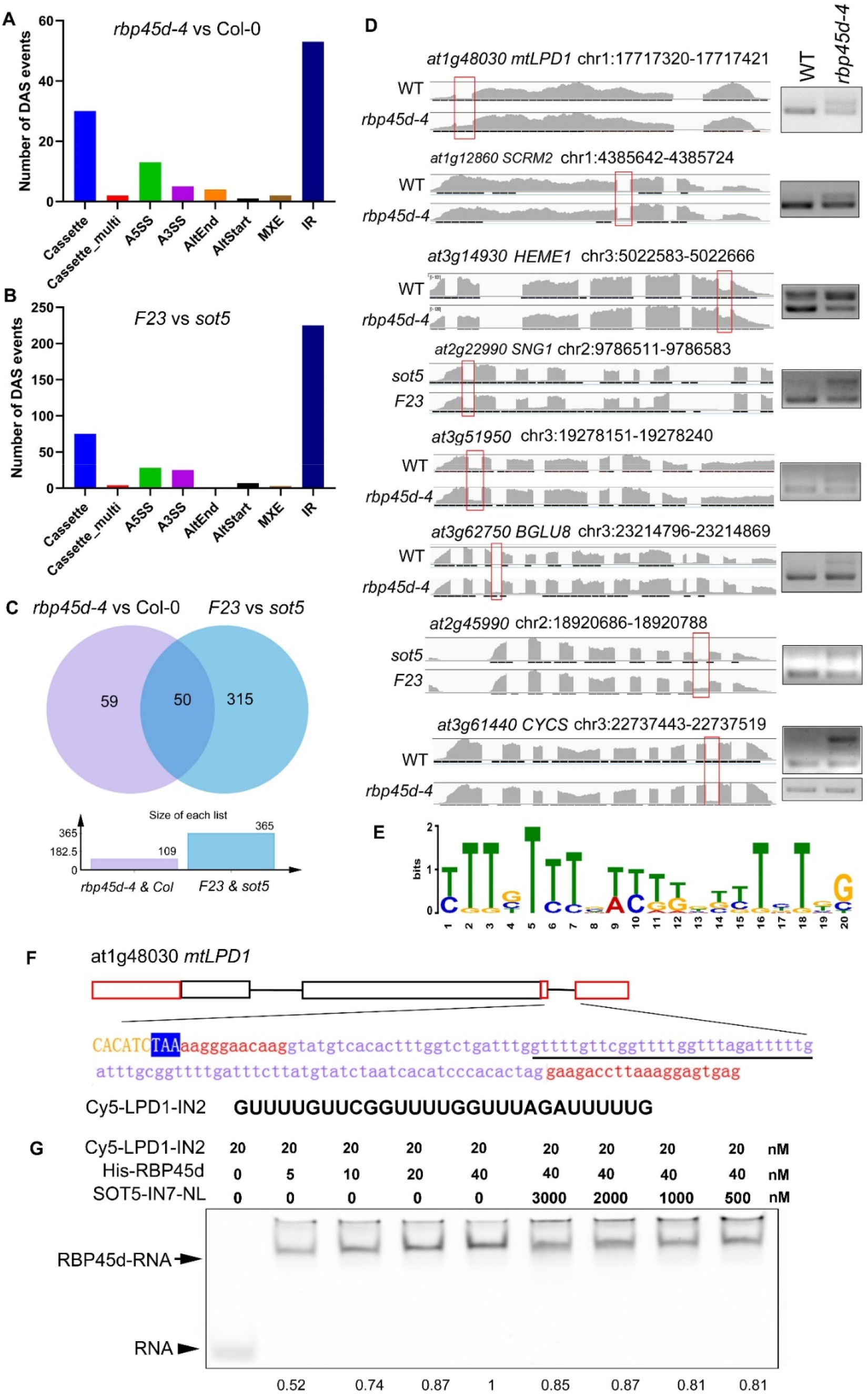
Effects of RBP45d mutations on splicing events and identification of U-rich sequences downstream of the 5’ ss. (A) and (B) Transcriptomic analysis of differential alternative splicing (DAS) events (FDR < 0.05) between WT and *rbp45d-4* and between *sot5* and *F23*, respectively. (C) Venn diagrams showing the number of unique or shared DAS in *rbp45d-4* and *F23*. (D) Representative Intron Retention (IR) events visualized by the Integrative Genomics Viewer browser (IGV) and validation by RT-PCR analysis between *rbp45d-4* and WT. The left panels show the coverage plots of each gene generated by IGV browser and the red rectangles marked the retained introns in *rbp45d-4* or *F23.* The right panels show RT-PCR analysis of the corresponding introns. The gel under the *CYCS* gene is *ACTIN2* which is used as an internal control. Two biological replicates (independent pools of aerial parts of plants) were analyzed, and one representative result is shown. (E) The T rich consensus motif is found from retained intron sequences by the MEME suite analysis. (F) The schematic presentation of the *mtLPD1* gene and its intron 2 sequence. The black boxes indicate the exons in CDS, the red boxes indicate the exons in UTR and the black lines indicate introns. The underlined letters in the intron correspond to the sequence of the RNA probe used in REMSA. (G) REMSA results show RBP45d protein can directly bind to the LPD1-IN2 probe. The unlabeled SOT5-IN7-NL probe can only partially compete with probe LPD1-IN2. The number at the bottom of each lane show relative levels of the RBP45d-RNA complex. The arrow showed the protein-RNA complex and the arrowhead showed the free RNA probe.

### RBP45d also modulates cryptic splicing site selection in the *clpR4-3* mutant

Previously, we reported the G to A mutation at the 5’ ss (+1G-to-A) of *ClpR4* intron 2 in the *clpR4-3* mutant, which displays virescent leaves (Wu et al., 2013). Similarly, the mutation of the authentic splice site leads to retention of intron 2 and activation of the two cryptic splice sites (+15 ss and +48 ss) (Figure 8A), producing three different transcripts, named band a, b, and c that are not detected in WT (Figure 8C). To examine whether RBP45d is involved in regulation of alternative splicing of *ClpR4* intron 2 in *clpR4-3*, we constructed the *rbp45d-4 clpR4-3* double mutant by genetic crossing. Although *rbp45d-4* was unable to suppress the virescent phenotype of *clpR4-3* (Figure 8B), the splicing site and splicing efficiency of *ClpR4* mRNA were clearly alerted in the double mutant, where band b products significantly decreased while band d products appeared and band c products increased slightly, compared with those in the single *clpR4-3* mutant (Figure 8C and 8d). Sequencing of the cloned PCR products showed that band b, c and d products were derived from the cryptic splice sites +48 ss, +15 ss, and −96 ss, respectively (Figure 8A, 8C and Figure S8). These results suggest that the absence of RBP45d in *clpR4-3* allows the U1 snRNP complex to preferentially recognize the cryptic splice sites −95 ss and +15 ss, instead of the cryptic splice site +48 ss. We found a five consecutive U (U-rich) element downstream of the cryptic splice site +48 ss. It is possible that *RBP45d* binds to this U-rich element in *ClpR4* intron 2, and subsequently promotes the cryptic +48 ss selection in *clpR4-3*.

**Figure 8.**
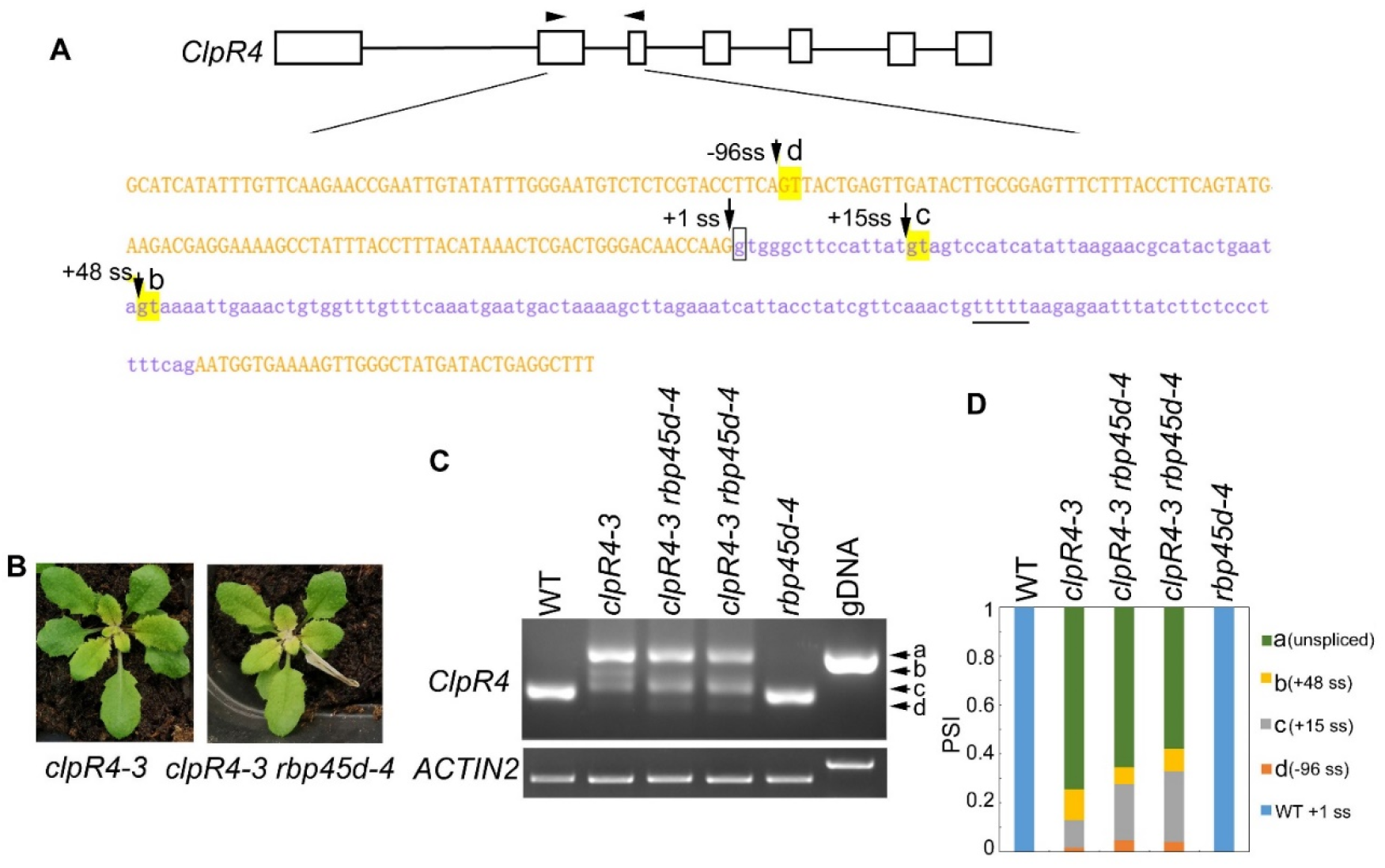
RBP45d regulates the cryptic 5’ splice site selection and splicing efficiency of *ClpR4* intron 2 in *clpR4-3*. (A) Schematic diagrams of the *ClpR4* gene and sequences of *ClpR4* exon 2 and intron 2. The black boxes indicate exons and the black lines indicate introns. Upper and lower case letters indicate exon and intron sequences, respectively. The arrows labelled with d, c and b indicate the activated cryptic 5’ splice sites at the −96 ss, +15 ss, and +48 ss, respectively. The arrow labelled with +1 ss indicates the authentic splice site, and boxed g indicates the point mutation in *clpR4-3*. The arrowheads indicate the primers used for RT-PCR. The T-rich sequence downstream the splice site +48ss is underlined. (B) Phenotype of *clpR4-3* and *clpR4-3 rbp45d-4* mutants. (C) Splicing patterns of *ClpR4* intron 2 in WT, *clpR4-3* and *clpR4-3 rbp45d-4* plants. Band a corresponds to the transcript retained intron 2, whereas band b, c and d correspond to the transcripts spliced at +48 ss, +15 ss and −96 ss, respectively. *ACTIN2* is used as an internal control. Two biological replicates (independent pools of aerial parts of plants) were analyzed, and one representative result is shown. (D) Relative levels of the PCR product bands in (C) quantified by ImageJ software. PSI (percent spliced in index) indicates the percentage of each variant in total transcripts.

### Mutations in *PRP39a* suppress the *sot5* phenotype

To clone the suppressor gene in the *E25* line, which displays a similar phenotype to *F23, F11* and *E1* but was not allelic with them, we used the backcrossed F_2_ population between *E25* and *sot5* to clone the suppressor gene via the MutMap approach. Our data showed that the suppressor gene was located in a region (~1.3 M) containing 15 candidate genes on Chromosome 1 (Figure S9A and S9B). Interestingly, we found a G to A mutation at the 5’ ss of the eleventh intron in the *PRP39a* gene, which encodes a U1 snRNP component PRP39a (Figure 9A). To verify the result, we identified a T-DNA insertion mutant *prp39a* (*prp39a-1*, salk_133733) from the ABRC stock, and made the *sot5 prp39a-1* double mutant by genetic crossing. The double mutant exhibited the same phenotype as *prp39a-1* (Figure 9B). Likewise, PCR analysis showed that the pattern of cryptic splicing products in *sot5 prp39a* was similar to that in *E25* (Figure 9C and 9D). In addition, we overexpressed the *PRP39a* genomic DNA in *E25*, and found that the phenotype and cryptic splicing pattern of the transgenic lines were the same as those of *sot5* (Figure 9C and 9D). Taken together, our results suggest that *PRP39a* mutations contribute to the suppression of *sot5* in *E25*.

**Figure 9.**
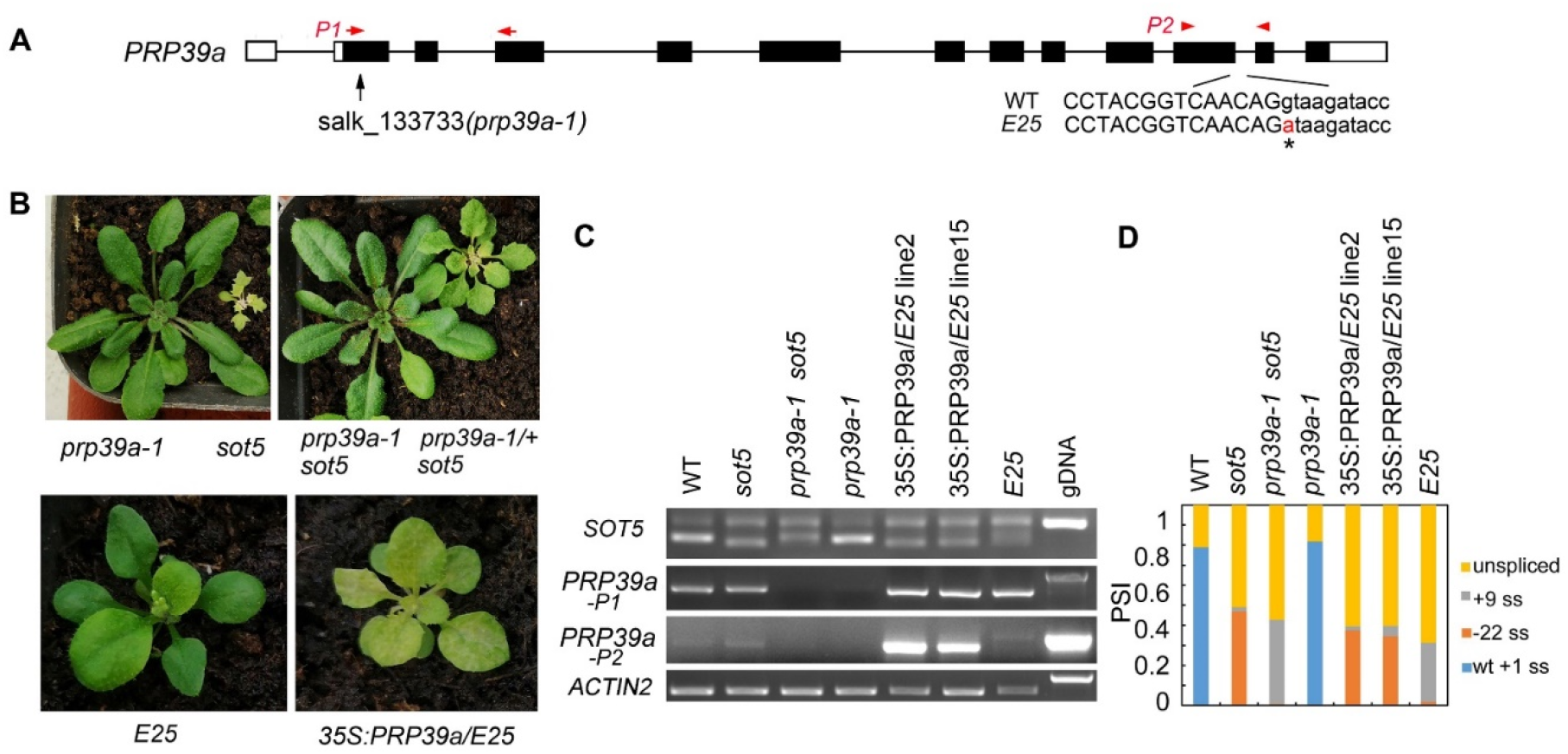
*PRP39a* is a new suppressor gene isolated from *E25*. (A) The schematic presentation of the *PRP39a* gene. The black boxes indicate exons and the black lines indicate introns. Black arrow showed the site of T-DNA insertion. The G to A point mutation of *E25* was indicated by *. And red arrow and arrowhead showed the primer pair P1 and P2 used for RT-PCR, respectively. (B) Upper panel, phenotypes of *prp39a-1, sot5, sot5 prp39a-1* and *PRP39a-1/^+^ sot5*. Lower panel, phenotypes of *E25* and *35S:PRP39a-gDNA/E25* transgenic plants. (C) RT-PCR analysis of the *PRP39a-1, sot5 prp39a-1* and *35S:PRP39a-gDNA/E25* transgenic plants. *ACTIN2* is used as an internal control. Two biological replicates (independent pools of aerial parts of plants) were analyzed, and one representative result is shown. (D) Quantification of the *SOT5* PCR product bands in (C). PSI (percent spliced in index) indicates the percentage of each variant in total transcripts.

### RBP45d specifically interacts with PRP39a

Our genetic screening of *sot5* suppressors indicated that RBP45d and PRP39a act in the same genetic pathway with regard to *SOT5* intron 7 alternative splicing. In addition, the *prp39a-1* mutant exhibited the same shorter primary root phenotype as *rbp45d*, compared to WT (Figure S11). We then examined whether PRP39a and RBP45d physically interact each other *in vitro* and *in vivo*. Y2H results showed that PRP39a directly interacted with RBP45d, but not with other RBP45/47 members (Figure 10A). Consistently, BiFC analysis showed that PRP39a and RBP45d interacted specifically in the nucleus (Figure 10B), which is consistent with the nucleus localization of PRP39a (Figure S10). However, we did not detect the fluorescence signal between PRP39a and RBP47b (Figure 10B), indicating that they are not associated each other *in vivo*. These results are consistent with previously reported that the role of RBP45d in alternative splicing relies on specific interaction with PRP39a (Chang et al, 2022). Meanwhile both Y2H and BiFC results showed that PRP39a was able to interact with U1C in the nucleus (Figure 10). Taken together, our results suggest that Arabidopsis RBP45d, PRP39a and U1C function in a similar way to that in yeast with regard to alternative splicing regulation.

**Figure 10.**
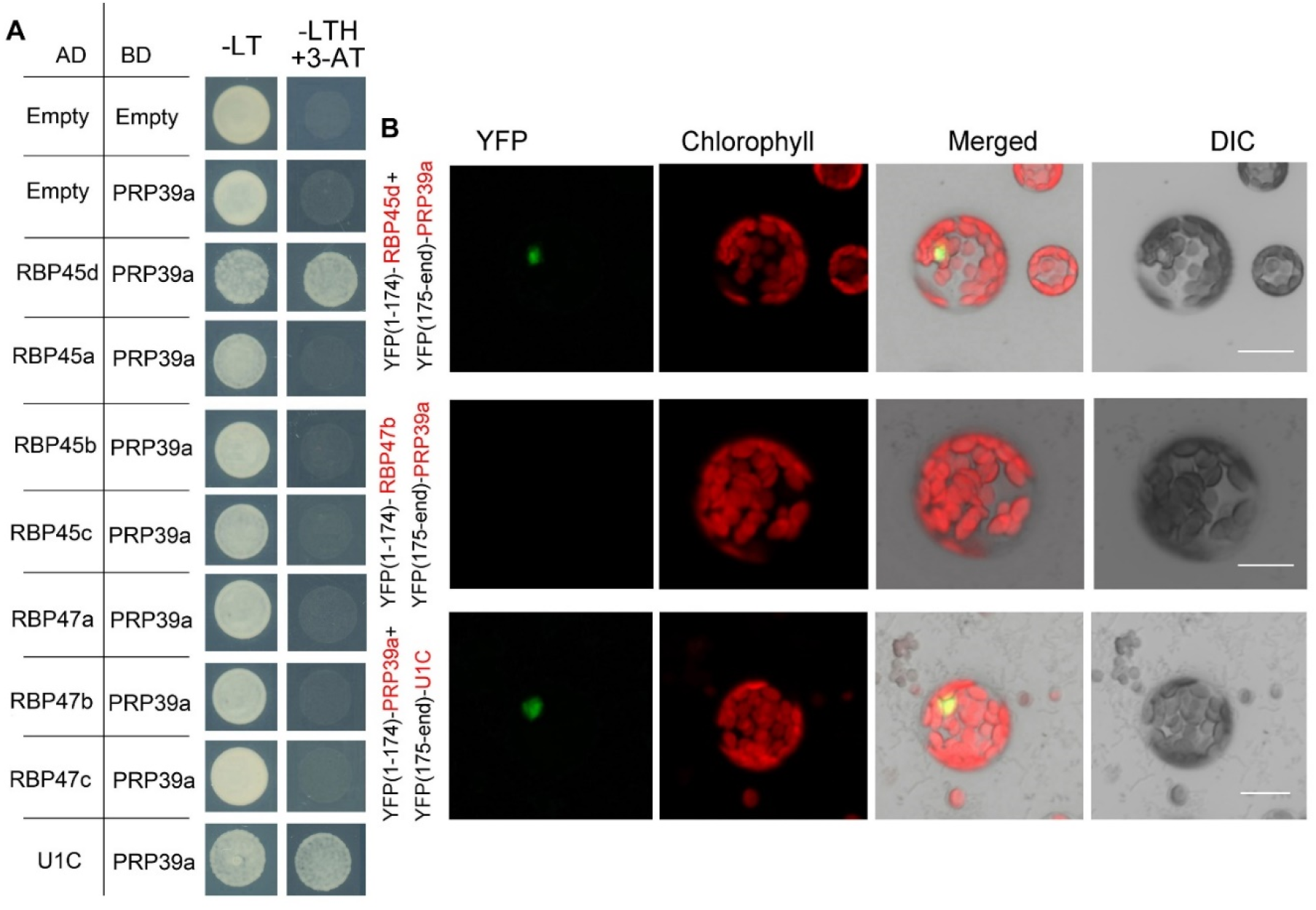
PRP39a interacts with RBP45d and U1C. (A) Yeast two-hybrid assays of the interaction between PRP39a and RBP45/47 members or U1C. The transformed yeast cells were grown on −LT (SD-Leu/-Trp) mediums and −LTH+3-AT (SD-Leu/ −Trp /-His with 5 mM 3-AT) mediums. (B) BiFC assay of the interaction between PRP39a and RBP45d, RBP47b or U1C. YFP, fluorescence of yellow fluorescent protein; DIC, Differential interference contrast microscopy; Merge, merge of YFP, DIC and chlorophyll. Bar = 15 μm.

## Discussion

In this study, we used a powerful genetic approach to screen phenotypic suppressors of the *sot5* mutant, which contains a G to A mutation at the first nucleotide of intron 7 (Huang et al., 2018), and isolated two suppressor genes, *RBP45d* and *PRP39a*. RBP45d and PRP39a are yeast Nam8 and Prp39 homologs, respectively, which are auxiliary components of the U1 snRNP complex. Although they have been demonstrated to play important roles in constitutive or alternative splicing in yeast, human and Arabidopsis (Puig et al., 1999, Le Guiner et al., 2001; Forch et al., 2002, Chang et al., 2022), their function in cryptic splicing remains unclear. Here we mainly characterized the function of RBP45d and collected several lines of evidence substantially supporting that RBP45d is a *trans*-acting splicing factor that bind to the U-rich element downstream the 5’ ss and subsequently facilitates spliceosome assembly. First, loss-of-function mutations in *RBP45d* lead to intron retention of a set of genes and significantly alter cryptic splicing frequency in the *sot5* mutant. Second, RBP45d directly interacts with the core subunit U1C of U1 snRNP and the U1 auxiliary factor PRP39a. Third, RBP45d can bind to U-rich sequences and its binding affinity is positively correlated to the U content of the sequence. Taken together, our data suggest that RBP45d together with PRP39a promotes 5’ ss selection by directly binding to U-rich *cis*-elements and then recruits U1 snRNP to the splice site (Figure 11).

**Figure 11.**
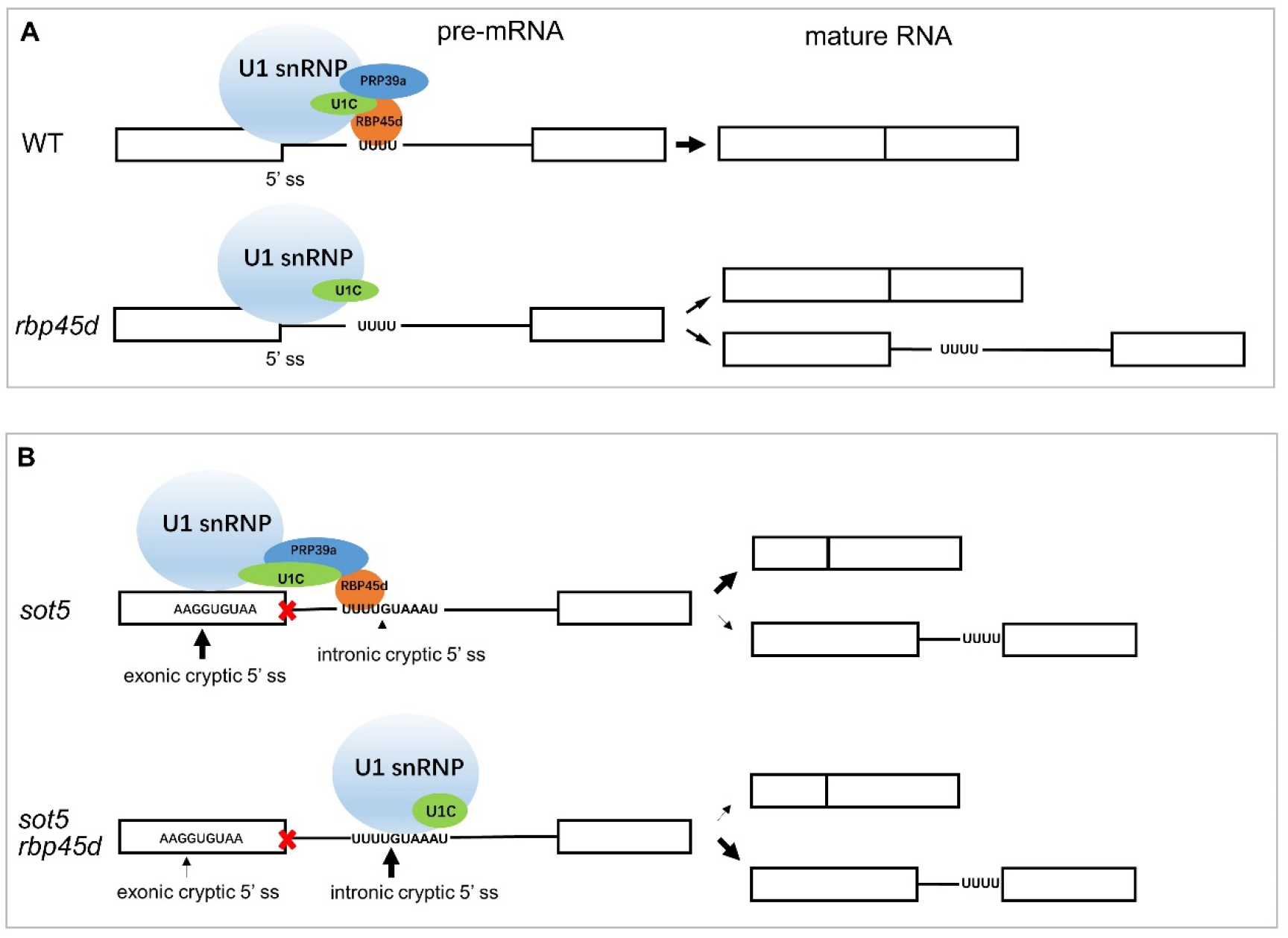
Working model of RBP45d in intron splicing and 5’ cryptic splice site selection. (A) In WT, RBP45d promotes the splicing efficiency of a set of U-rich introns such as *mtLPD1* intron2 by binding to the intronic U-rich element. Meanwhile RBP45d interacts with PRP39a and U1C so as to stabilize the U1 snRNP at the 5’ ss and then recruit spliceosome. When RBP45d is absent, the splicing efficiency of the U-rich intron significant decreased probably due to the less or unstable U1 snRNP (lacking PRP39a and RBP45d) formed at the 5’ ss. For those alternative splicing intron, RBP45d is a splicing enhancer. (B) In the 5’ ss intron mutant, for example, *sot5*, the authentic 5’ ss no longer base paired with U1 snRNA or base paired with U1 snRNA imperfectly, so that the exonic and intronic cryptic 5’ ss nearby are activated. RBP45d binds to the U-rich element located downstream of the exonic cryptic 5’ ss and then promote U1 snRNP to select the 5’ ss. It is also likely that the RBP45d bound to the U-rich element which is proximal to the intronic cryptic 5’ ss mask the 5’ ss. When RBP45d is absent, the masked intronic cryptic 5’ ss is exposed and preferred by the U1 snRNP complex (lacking PRP39a and RBP45d). The black boxes indicate exons and blank lines indicate introns. UUUU indicates U-rich element. The nine nucleotide AAGGUGUAA is the exonic cryptic 5’ ss in *sot5* intron7. The nine nucleotide UUUGUAAAU is the intronic cryptic 5’ ss in *sot5* intron7. The red crosses indicate 5’ ss mutation. The black arrows indicate the usage of the 5’ ss and the thickness of the arrows indicate the frequency of the splice site usage.

### RBP45d regulates alternative splicing

It is well known that U1 snRNP plays a critical role in initiation of 5’ ss recognition, and several U1 snRNP auxiliary factors can influence 5’ ss selection (Puig et al., 1999; Fortes et al.,1999; Forch et al., 2002). Based on the data derived from this study and reported previously, the U1 snRNP component RBP45d is required for alternative splicing for a set of genes in plants (Chang et al., 2022). Consistently, the two RBP45d homologs, yeast Nam8 and human TIA-1, are implicated in alternative splicing via binding to the U-rich region downstream of a weak 5’ ss (Puig et al., 1999, Qiu et al, 2011, Le Guiner et al., 2001, Aznarez, et al, 2008). The same biological function of Nam8, TIA-1, and RBP45d is attributed to their highly conserved domains including the three RRM domains and the C-terminal Gln-rich tail. Cryo electron microscopy (cryo-EM) analysis of prespliceosome structure revealed that Nam8 or TIA-1 bind to U-rich sequences through the RRM2 domain to define the use of weak 5’ ss, and interacts with the U1C-terminal region near the U1-5’ ss helix via the RRM3 domain (Plaschka et al., 2018). Recently, Cryo-EM structure of U1 snRNP showed that the alternative splicing factors Prp39 and Prp42 form heterodimer that acts as a central scaffold connecting all auxiliary proteins associated with U1 snRNP to the core (Li et al., 2017). In human, there is no Prp42 homolog, but the Prp39 homolog PrpF39 forms a homodimer that acts as the Prp39/Prp42 heterodimer in U1 snRNP. In addition, Nam8/TIA-1 plays a role in the U1-U2 snRNP complex formation through interaction of the RRM3 domain and C-terminal tail with the Prp39/Prp42 heterodimer (Plaschka et al., 2018). The structural and biochemical data provide insight into alternative splicing mediated by Nam8/TIA-1 and Prp39/prp42. Although the U1 snRNP complex and structure remain to be dissected in plants, most of the counterparts of yeast U1 snRNP core subunits and auxiliary proteins are present in Arabidopsis and encoded by multiple genes (Wang and Brendel, 2004). Considering that the basic mechanism of pre-mRNA splicing is highly conserved in all eukaryotic organisms, we propose that RBP45d functions as a splicing factor in the same way as Nam8/TIA-1. As expectedly, our data showed that RBP45d can not only bind to U1C (the core protein of U1 snRNP) and PRP39a (the scaffold protein connecting the auxiliary proteins to the core), but also to the downstream U-rich RNA sequence near 5’ ss of *SOT5* intron 7 and *mtLPD1* intron 2. These results suggest that RBP45d can recruit spliceosome to specific 5’ ss and enhance the stability of spliceosome by directly binding to U-rich RNA sequences downstream of 5’ ss.

### RBP45d regulates cryptic splicing in 5’ ss mutants

To date, it remains unknown whether the alternative splicing factor Nam8/TIA-1/RBP45 is also involved in cryptic splice site selection. In our model system, the mutation at the authentic 5’ ss of *SOT5* intron7 leads to activation of two cryptic splice sites at which the +9 ss is much weaker than the −22 ss (Figure 1B). Polypeptide prediction showed that mRNA derived from +9 ss encodes a protein with functionally similar to but three amino acid residues more than endogenous SOT5, while that from −22 ss generates a truncated protein without function. The *sot5* mutant exhibits a leaf virescent phenotype due to a certain level of functional SOT5 still produced, otherwise is embryonic lethality. However, RBP45d mutations significantly reverse the splicing efficiency of the two cryptic splice sites, which well explains why null mutations of *RBP45d* can completely rescue the *sot5* phenotype to the WT level. Careful inspection of the sequence nearby the cryptic splice sites showed a U-rich element located downstream of the −22 ss but not of +9 ss. REMSA experiments indeed demonstrated that RBP45d can directly bind to this U-rich fragment, and their binding affinity gradually decreases with the decrease of U and A content *in vitro*. These results suggest that RBP45d plays a role in selection of cryptic splice sites upstream of the RBP45d-bound U-rich element. To testify whether RBP45d can also regulate cryptic splice sites selection in other mutants, we constructed the double mutant of *clpr4-3 rbp45d*. *clpr4-3* has a G to A mutation at the first nucleotide of intron 2. Similarly, the authentic splice site mutation in *clpr4-3* results in activation of two cryptic splicing sites at position +15 ss and +48 ss. Interestingly, *RBP45d* mutations also obviously changed cryptic splicing patterns of *clpR4-3* pre-mRNA, accompanying with a significant decrease in the +48 ss usage. We searched for U-rich elements nearby the cryptic splice sites and found a U-rich sequence only downstream of the +48 ss, which was probably bound by RBP45d.

Cryptic splicing is repressed by complicated mechanisms in eukaryotes. In human disease, cryptic splice sites can be activated by different mutations including in *cis*-acting silencing elements, *trans*-acting repressors or the core splicing machinery (Annal and Monika 2018). Recently, a *Caenorhabditis elegans* genetic screen identified that mutations in Prp8, a core spliceosome component, alter cryptic splicing frequency, but almost have no effect on alternative splicing patterns, suggesting that the mechanism regulating cryptic splicing is distinct from that of alternative splicing (Mayerle et al, 2019). Spliceosome has evolved intrinsic ability to prevent from uses of cryptic splice sites in pre-mRNA splicing. Interestingly, targeted RNA-seq analysis using a large library of synthetic introns in yeast revealed a similar mechanism by which splicing machinery can prefer to utilize alternative 3’ ss downstream of the original site, thus avoiding selection of cryptic splice sites nearby the 3’ end of introns (Schirman, et al, 2021). Further analysis showed that the vast majority of cryptic splicing sites located downstream of 3’ss sites are activated by a loss of a specific splicing factor U2AF35 (Schirman, et al, 2021). Taken together, many *trans*-acting splicing factors are involved to prevent occurrences of cryptic splicing.

Increasing evidence suggests that spliceosome has been evolved to distinguish authentic splice sites from cryptic and pseudo splice sites (Roca et al., 2003; Dawes et al., 2022). However, cryptic splicing of pre-mRNA is inevitable since mutations in either *cis*-acting elements and/or *trans*-acting factors take place *in vivo* frequently. It is possible that mRNA variants derived from cryptic splice sites nearby the authentic splice site can generate a partially functional protein or even a full functional protein, which alleviates the effect caused by the mutations. In fact, many mutants with mutations at splice sites such as *sot5* and *clpr4-3* have been reported to exhibit a weaker phenotype than their corresponding null mutants. Thus, from the viewpoint of evolution, activated cryptic splice sites might become a driven source in developing new alternative splice sites, since such mutants are survival and capable to reproduce.

### The distinctive function of RBP45d from other members in pre-mRNA splicing

There are eight paralogs in the Arabidopsis RBP45/47 family. Whether they function redundantly or not remains unclear. Here we showed that among these RBP45/47 members RBP45d can specifically allow U1 snRNP to bind to 5’ ss. The specificity of RBP45d in pre-mRNA splicing is supported by the following evidence. The *sot5* phenotype can be rescued by mutations in *RBP45d* but not in other *RBP45/47* members. Only RBP45d is interacted with PRP39a, since other RBP45/47 members lack the long C-terminal tail that was demonstrated to play an important role in protein-protein interaction between RBP45d and PRP39a (Chang et al., 2022). Furthermore, we observed that *rbp45d* mutants have short root and late flowering phenotypes, which are not present in other *rbp45/47* mutants (Figure 4 and Figure S7). However, we found some common features in all the RBP45/47 members. For example, they are localized in both the nucleus and cytoplasm (Figure S5), and interact with the U1 snRNP core subunit U1C (Figure S6). Thus, it is likely that other RBP45/47 members are also involved in splicing regulation through the mechanisms different from those of RBP45d. It will be interesting to investigate biological functions of other RBP45/47 members as well as the molecular mechanisms by which they act in plants.

## Materials and Methods

### Plant materials and growth conditions

The Arabidopsis ecotype Columbia-0 (Col-0) was used as the wild type (WT) in this study. The mutants *sot5* and *clpR4-3* were previously described (Huang et al., 2018, Wu et al., 2014). The *prp39a-1* (salk_133733), *rbp45a* (salk_140650), *rbp45c* (salk_063484), *rbp47a* (salk_142402), and *rbp47b* (GK-626C01) mutants were obtained from the ABRC stock center (https://abrc.osu.edu/) and were genotyped by PCR using gene specific primers (Table S7). Double mutants *prp39a-1 sot5, rbp45a sot5, rbp45c sot5 rbp47a sot5 rbp47b sot5,* and *clpR4-3 rbp45d-4* were identified from F_2_ generations derived from crosses between single mutants by the PCR-based genotyping procedure. Seeds were surface-sterilized by 75% ethanol and stratified at 4°C for 3 days, and then sown onto half-strength Murashige and Skoog (MS) medium with 0.5% sucrose. Seeds were also directly sown in soil and grown in a phytotron with long-day conditions (16 h light/8 h dark) and light intensity (100 μmol photons m^-2^ s^-1^) at 22°C.

### EMS mutagenesis, gene cloning, plasmid construction and transformation

In order to obtain *sot5* suppressors, *sot5* seeds were ethyl methanesulfonate (EMS) mutagenized (0.1%). About 30000 M_2_ plants from 5000 M_1_ lines were screened. Several suppressor lines that displayed WT-like green leaves were obtained from the M_2_ generation. To clone *F23*, we produced an F_2_ population from the cross between *F23* and Landsberg *erecta* (L*er*). In the F_2_ population, about eighty *F23* plants were pooled for Whole Genome Resequencing and Mapping-By-Sequencing (MBS) analysis (OE Biotech Co.,Ltd). To clone the *E25* gene, we crossed the *E25* with *sot5* and generated the BC_1_ F_2_ population. About thirty *E25* plants were pooled for Whole Genome Resequencing and Mutmap analysis (Biomarker Technologies).

The *RBP45d* coding DNA sequence (CDS) and *PRP39a* genomic DNA were cloned into the pENTR SD/D-TOPO entry vector (Invitrogen), and then recombined into the pGWB2 destination vector, respectively. The destination vector containing *RBP45d* was transformed into *F23* mutant and the destination vector containing *PRP39a* was transformed into *E25* mutant according to the method as our previously described (Huang et al., 2018). To obtain recombinant RBP45d protein, the *RBP45d* CDS was transferred into the pet 51b vector and then was fused with His tag at its N-terminal.

### Generation of *rbp45d-CR* mutant line by CRISPR/Cas9-mediated genome editing

The *rbp45d-CR* mutant line was generated by CRISPR/Cas9-mediated genome editing. The sgRNA sequences were designed by the online software CRISPR-P v2.0 (Liu et al., 2017) and CRISPR-GE (Xie et al., 2017) to target exon 2 of the *RBP45d* gene. The binary vector pYAO-Cas9-1300 (Feng et al., 2018) was linearized using *Bsa*I (ThermoFisher Scientific, UK) endonuclease. Two complementary sgRNA nucleotides were synthesized and annealed to generate double-strand DNA with appropriate overhangs on both ends and then inserted into the pYAO-Cas9-1300 vector. The vector was transformed into the *Agrobacterium tumefaciens* strain GV3101 competence cells. And the GV3101 with the vector was transformed into the Col-0 wild-type *Arabidopsis* genome by the floral dip method. The transgenic plants of T_1_ generation were screened by Hygromycin B resistance, DNA from positive lines was extracted and PCR and sequencing were performed to identify mutations in sgRNA target sequences. Finally, we screened the Cas9-free background and homozygous *RBP45d* gene editing lines in T_2_ generation plants to obtain *rbp45d-CR* mutant lines.

### Analysis of RT-PCR, RNA-seq and differential Alternative Splicing events (DAS)

For RT-PCR, total RNAs were extracted from seedlings or different plant tissues by RNA Easy Fast Plant Tissue Kit (TIANGEN, DP452) according to the manufacturer’s instructions. DNase I treatment and RT-PCR analysis were conducted as previously described (Huang et al., 2013). Semi-quantitative RT-PCR was carried out using the gene-specific primers (Table S7). To identify the splicing variants, the PCR products amplified by the primers spanning the introns were purified and cloned into pMD19-T vector (TaKaRa, D102A). About 10~20 clones of each PCR products were sequenced and aligned with the corresponding genes by SnapGene software.

Transcriptome sequencing and analysis were performed by NovelBio Company (http://www.novelbio.com/case/jycx/1.html). Total RNAs extracted with Trizol reagent (Invitrogen) from leaves of 25-day-old WT, *sot5*, *F23* and *rbp45d-4* seedlings were treated with Turbo RNase-free DNase I (Thermo Fisher Scientific) to remove genomic DNA. Polyadenylated RNA was isolated using magnetic beads with Oligo (dT). The RNA-seq libraries were constructed with the Illumina Whole Transcriptome Analysis Kit as the standard protocol described (Illumina, HiSeq system), and were sequenced on the HiSeq 2000 platform. The sequencing data were analyzed by NovelBio Company. To identify differential AS events, we subjected the RNA-seq data to analysis by CASH (Comprehensive AS Hunting) software (Wu et al., 2017). Statistical parameters for significant splicing change were defined as FDR < 0.05 and P-value < 0.05 (Table S2 and S3). The selected intron retention (IR) events were validated by RT-PCR using a set of primers (Table S7) that were designed based on each IR event.

### Subcellular localization of RBP 45/47 members and PRP39a fused with YFP and BiFC assay

To express YFP-fused protein, the full length CDS of the *RBP45a, RBP45b, RBP45c, RBP45d, RBP47a, RBP47b, RBP47c, RBP47c’, PRP39a* and *U1C* genes were amplified from the cDNA of WT seedlings using the primers (Table S7). The sequences were cloned into the pENTR SD/D-TOPO entry vector (Invitrogen) and then recombined into the p2YWG7 vector (http://gateway.psb.ugent.be/vector/show/p2GWY7/search/index/) or BiFC vctors pE3130 and pE3136 (http://www.bio.purdue.edu/people/faculty/gelvin/nsf/protocols_vectors.htm). The purified plasmid was transformed into the *Arabidopsis* protoplasts according to the method (Wu et al., 2009). The BiFC vector combinations are indicated in the Figure 5 and Figure 10.

### Yeast two hybrid

The CDS of *RBP45a, RBP45b, RBP45c, RBP45d, RBP47a, RBP47b, RBP47c, PRP39a U1A, U1C* and *U1-70K* were clone into the pENTR vector and then recombined into pDEST22 and pDEST32 destination vectors, respectively. For yeast two-hybrid assays, plasmids were transformed into yeast strain AH109 by the lithium chloride–polyethylene glycol method according to the manufacturer’s manual (Clontech). The transformants were selected on SD-Leu-Trp plates. The protein– protein interactions were tested on SD-Trp-Leu-His plates with or without 5mM 3-amino-1,2,4-triazole (3AT).

### Protein expression, purification and RNA electrophoretic mobility shift assay (REMSA)

The RBP45d CDS was cloned into the expression vector pet51b, and protein expression was induced using 0.5 mM isopropyl-b–D–thiogalactopyranoside at 16°C for 12 h in E. coli strain Rosetta. The protein was purified as described by Zhu et al., (2020). The RNA probes were chemically synthesized, and their 5’-end were labeled by Cy5 (Sangon Biotech (Shanghai) Co., Ltd). The sequences of the probes were as follows:

Cy5-SOT5-IN7: 5’-(CY5) CUCAAUAUGAUAAAUUUUGUAAAUC-3’;

Cy5-LPD1 IN2: 5’- (CY5) GUUUUGUUCGGUUUUGGUUUAGAUUUUUG-3’;

The corresponding non-labeled and mutated probes were also chemically synthesized for competition assays.

SOT5-IN7-NL: 5’-CUCAAUAUGAUAAAUUUUGUAAAUC-3’

SOT5-IN7-M1-NL: 5’-CUCAAUAUGGUAAAUUUUGUAAAUC-3’

SOT5-IN7-M2-NL: 5’-CUCAAUAUGGCAAAUUUUGUAAAUC-3’

SOT5-IN7-M3-NL: 5’-CUCAAUAUGGCAAACCGGGUAAAUC-3’

SOT5-IN7-M4-NL: 5’-CUCAAUAUGCCCGGCCGGGUAAAUC-3’

For EMSA assay, RBP45d protein and RNAs were incubated at room temperature for 30 min in the reaction solution (10 mM HEPES pH 7.3, 20 mM KCl, 1 mM MgCl2, 1 mM dithiothreitol, 5% glycerol (v/v) and 0.1 μg tRNA). The reactants were analyzed using a native polyacrylamide gel, and signals were detected by Azure Biosystems C600.

## Supplemental Tables

**Table S1.** The number of significantly differential AS events (DAS) in pairs of *rbp45d-4* vs. Col and *F23* vs. *sot5*.

**Table S2.** DAS events in the pair of *rbp45d-4* vs. Col.

**Table S3.** DAS events in the pair of *F23* vs. *sot5*.

**Table S4.** List of common DAS events of the two groups.

**Table S5.** Common DAS events of the two groups.

**Table S6.** The sequences of introns which were significantly retained in *rbp45d-4* and *F23*.

**Table S7.** Primers used in this study.

**Table S8.** Gene accession numbers in this study.

## Supporting information

Supplemental Data Sets

## ACKNOWLEDGMENTS

We thank the ABRC for providing the *prp39a-1* (salk_133733), *rbp45a* (salk_140650), *rbp45c* (salk_063484), *rbp47a* (salk_142402), and *rbp47b* (GK-626C01) seeds. We thank Yu Fang, Xiong Renxue and Wang Yanzhu for purifying the RBP45d protein. We thank Zhang hui and Wu Wenjuan for designing the sgRNA targeted *RBP45d*. This work was supported by the National Natural Science Foundation of China (32171293), The Fund of Innovation Program of Shanghai Municipal Education Commission (2021-01-07-00-02-E00117), and Shanghai Engineering Research Center of Plant Germplasm Resources (17DZ2252700).

## AUTHOR CONTRIBUTIONS

W.H. and J.H. designed the research; W.H., L.Z., YJ.Z., J.C., YW.Z., and F.L. performed the experiments; W.H. and J.H. supervised the experiments and wrote the article.

## Supplemental figures

**Figure S1.**
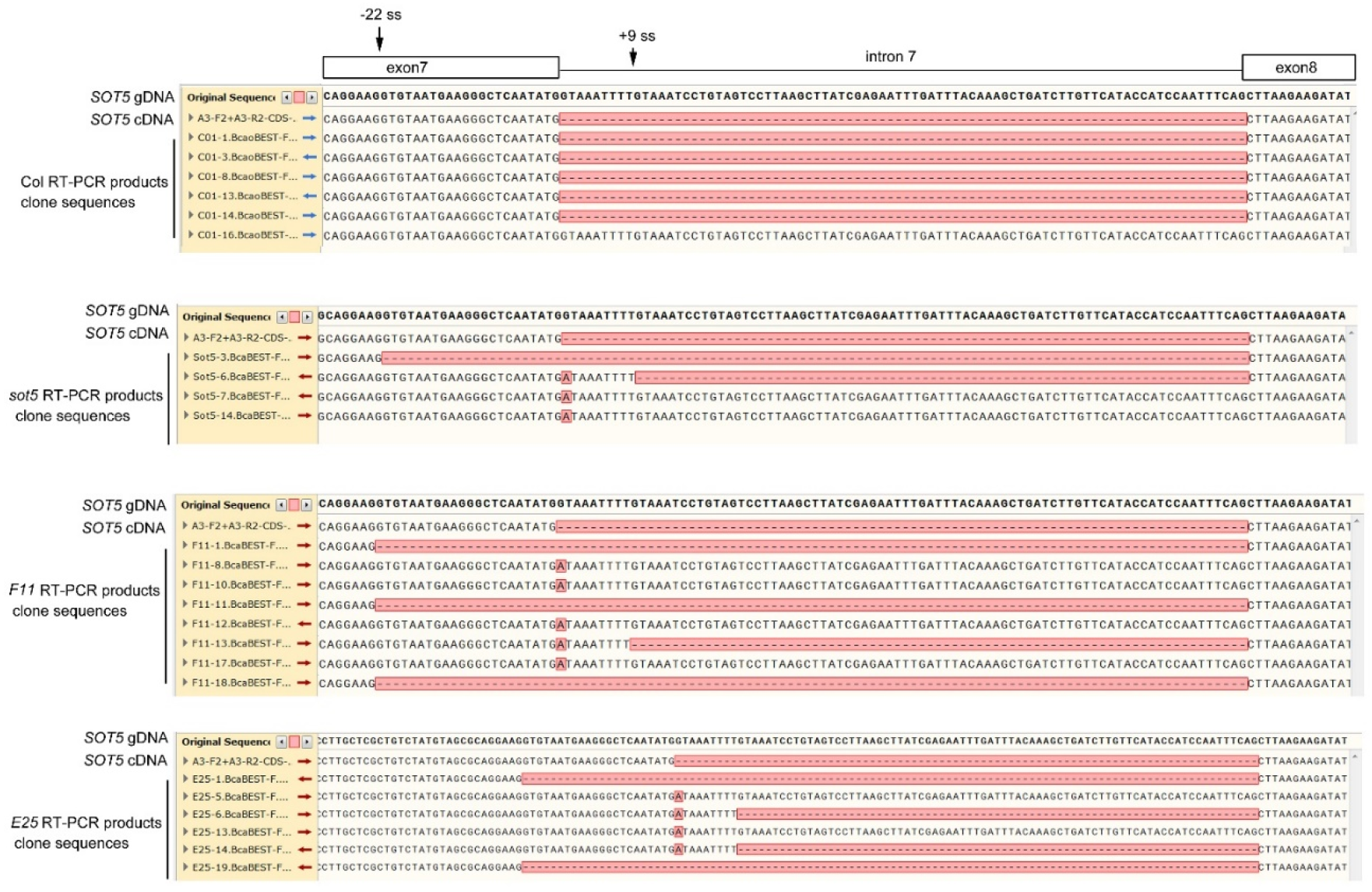
Sequence analysis of the cloned RT-PCR products amplified across *SOT5* intron 7 in WT, *sot5* and the suppressor lines *F11* and *E25*. The DNA sequences from different clones are aligned by SnapGene software. The sequences nearing *SOT5* intron 7 are showed.

**Figure S2.**
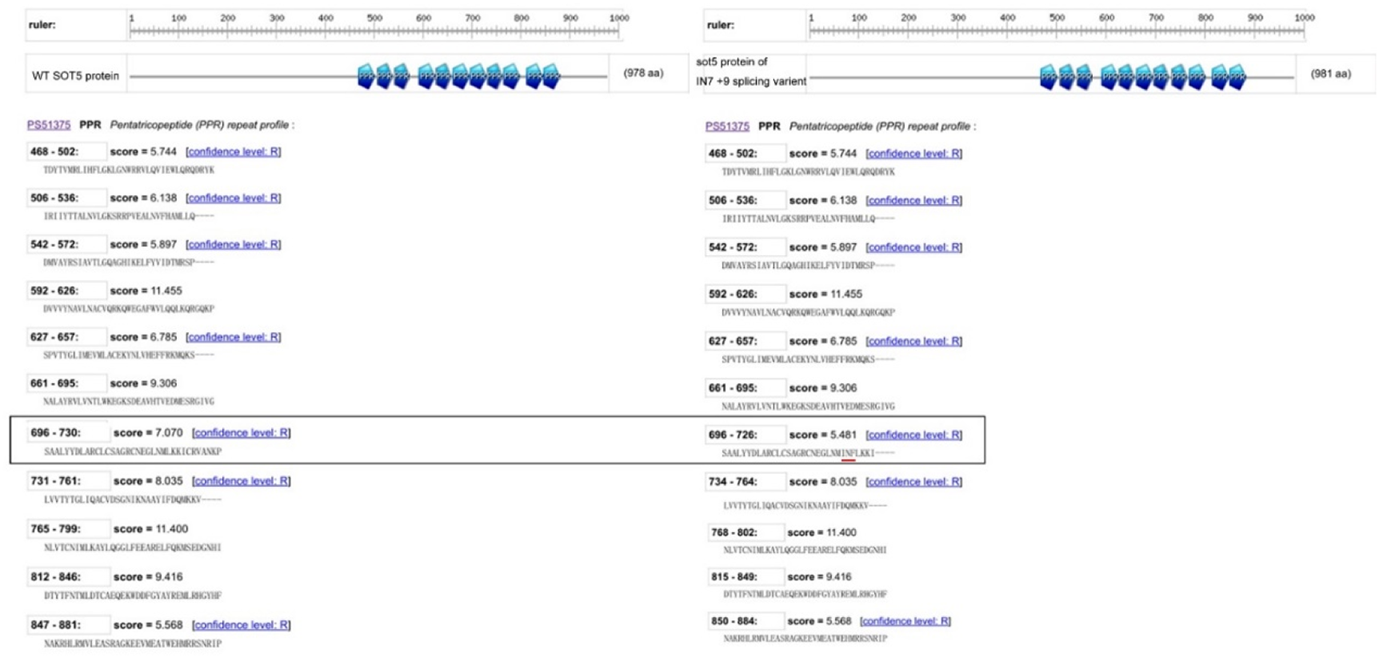
Comparison of the two SOT5 proteins encoded from wt +1 ss and +9 ss products. The domains of different protein variants were predicted by Prosite (https://prosite.expasy.org/). The 978 aa and 981 aa SOT5 proteins are generally the same as the endogenous one except that the seventh PPR domain (marking by the black box) is slightly altered.

**Figure S3.**
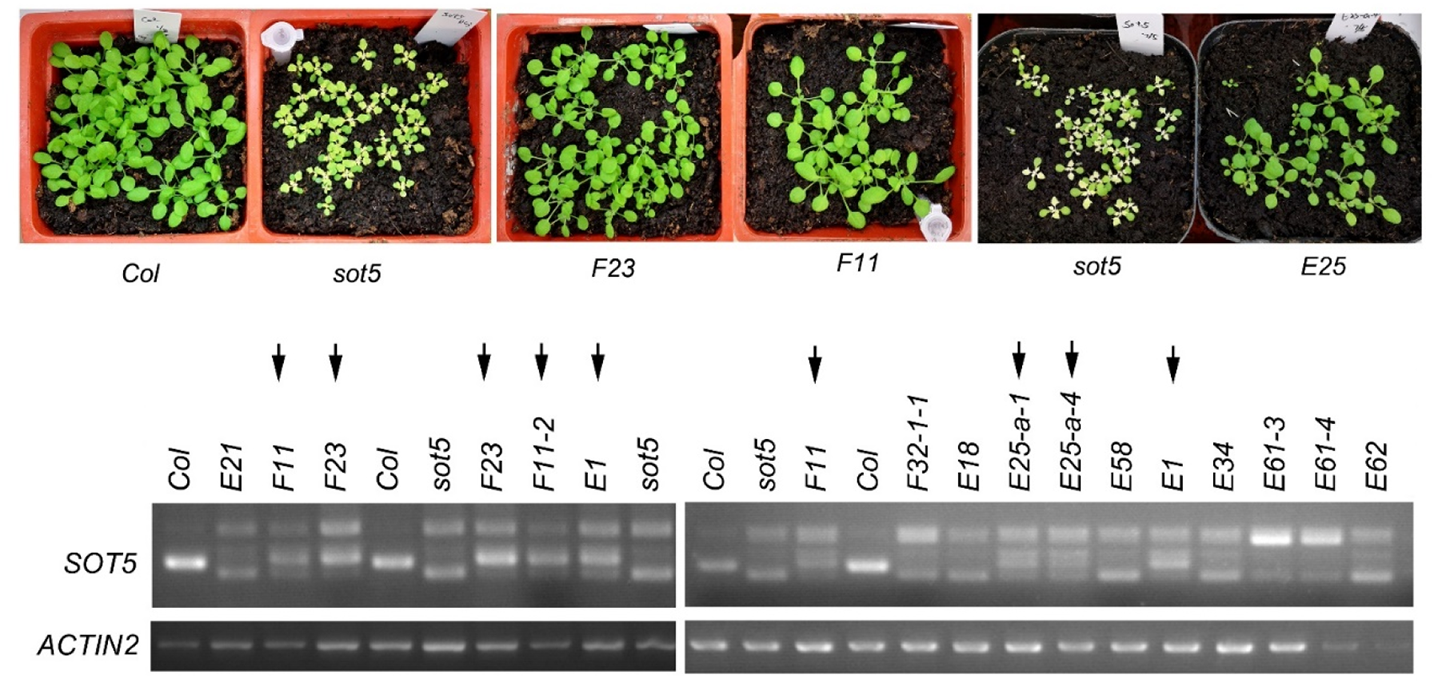
Phenotypes of *sot5* suppressors and their splicing patterns of *SOT5* intron 7. The upper panel shows the four suppressors lines presented the WT-like phenotype. The lower panel shows the splicing patterns of *SOT5* intron 7 in different suppressors lines. Among these suppressors, *F23, F11, E1* and *E25* (indicated by the arrows) show a similar splicing pattern of *SOT5* intron 7.

**Figure S4.**
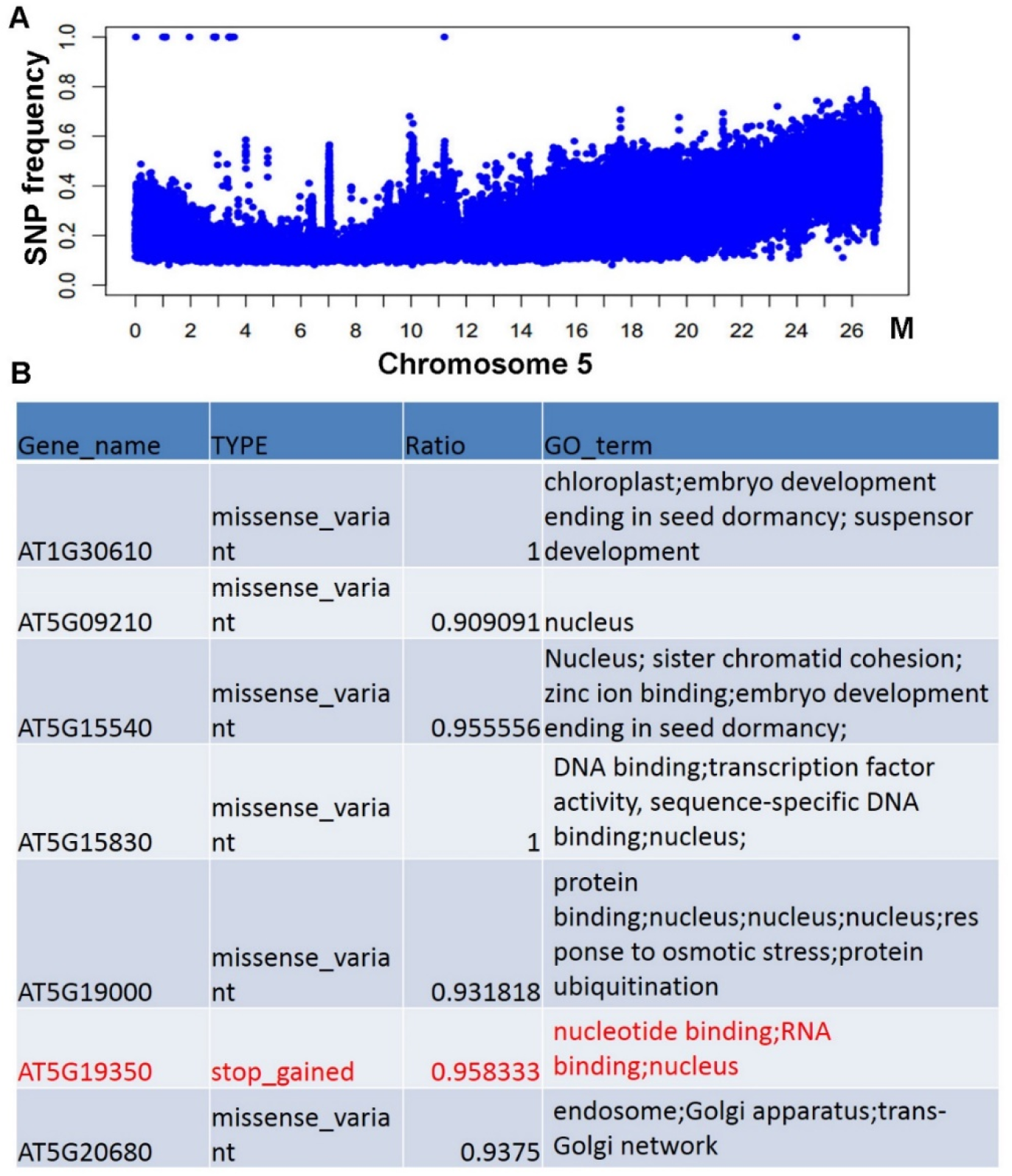
Cloning the *F23* suppressor gene through the Mapping-By-Sequencing (MBS) technique. *F23* (Col background) was crossed with L*er* to generate F_2_ population for gene mapping. About 80 WT-like plants with *sot5* mutation in F_2_ population were selected and pooled for DNA extraction and genome re-sequencing. The frequency of SNP polymorphisms on each chromosome was then analyzed. Theoretically, *F23* mutation is closely linked with Col type SNPs. The L*er* SNP frequency close to *F23* mutation is about 0. SNPs not linked to *F23* should be half Col type and half L*er* type. (A) The frequency of SNP on chromosome 5 where *F23* mutation is located, and the blue dots represent SNP molecular markers. (B) Suppressor gene in *F23* was mapped to the interval containing six candidate genes with missense mutations and one candidate gene with the premature stop codon. And the at5g19350 which encodes the pre-mRNA splicing factor RBP45d is the most likely candidate gene.

**Figure S5.**
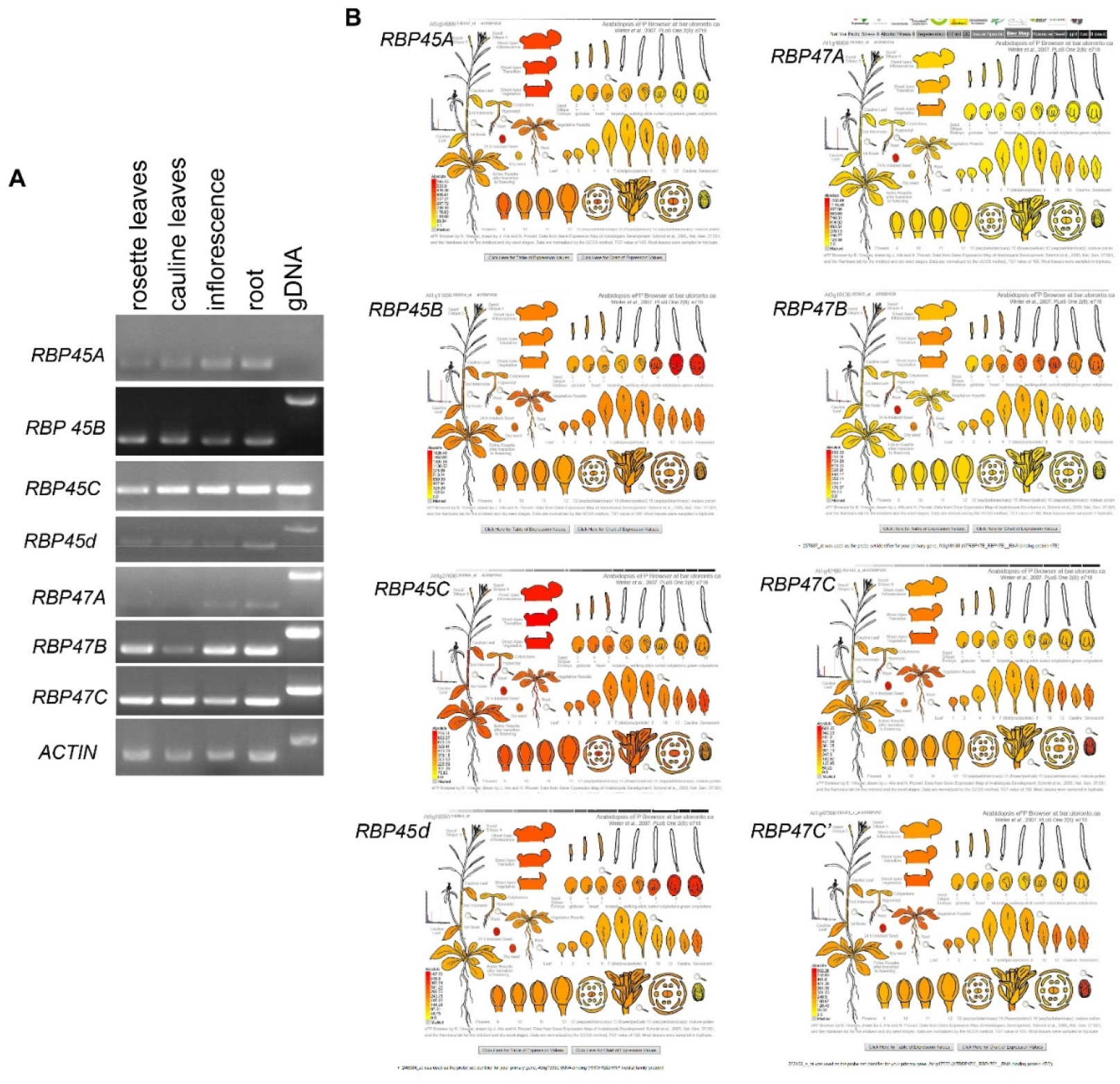
Expression patterns of *RBP45/47* family genes by RT-PCR and eFP browser. (A) RT-PCR analysis of *RBP45/47* gene expression in different organs. (B) eFP browser (http://bar.utoronto.ca/efp_arabidopsis/cgi-bin/efpWeb.cgi) shows expression patterns of *RBP45/47* genes in different organs.

**Figure S6.**
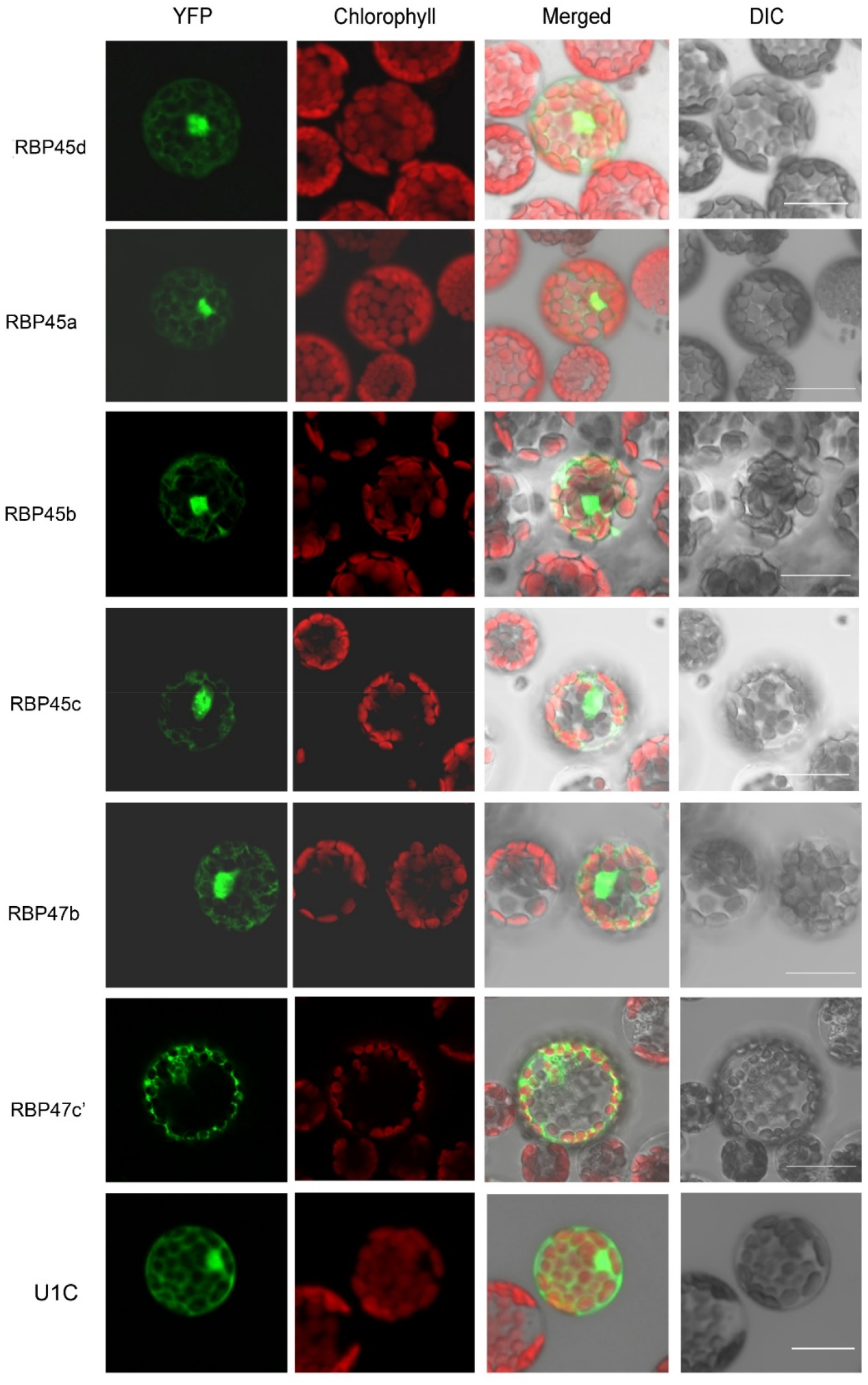
Subcellular localization of RBP45/47 family proteins and U1C. Confocal microscopy analysis of YFP fusion proteins transiently expressed in Arabidopsis protoplasts. The yellow fluorescence of the proteins was overlapped with chloroplast autofluorescence in merged images. DIC, Differential interference contrast microscopy. Bars = 15 μm

**Figure S7.**
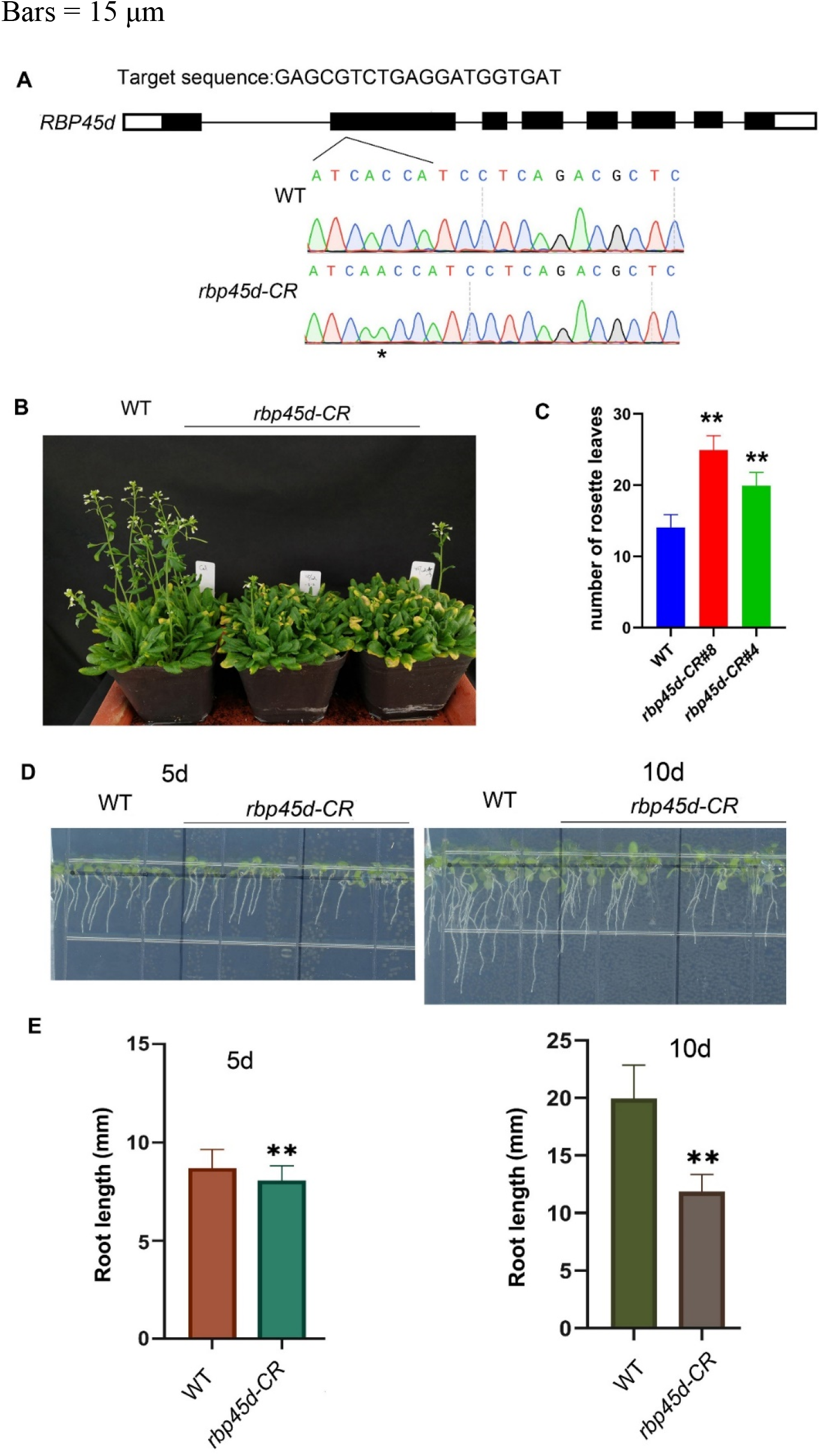
Phenotypes of *rbp45d-CR* plants generated by CRISP/CAS9 technique. (A) The sgRNA sequence and the mutation site of the *rbp45d-CR* plant. * indicates the inserted A which led to frameshift of RBP45d. (B) The phenotype of 45d-old *rbp45d-CR* plants grown under the 16h light/8h dark photoperiod. (C) The statistical analysis of *rbp45d-CR* rosette leaf number when flowering. The data are means ± SD (n = 20). Student’s *t*-test; **, *P* < 0.01. (D) Short primary root phenotype of the 5-d- and 10-d-old *RBP45d-CR* seedlings. (E) Quantification of root length shown in (D). The data shown are means ± SD (n = 20). Student’s *t*-test; **, *P* < 0.01. The experiments were performed at least two times.

**Figure S8.**
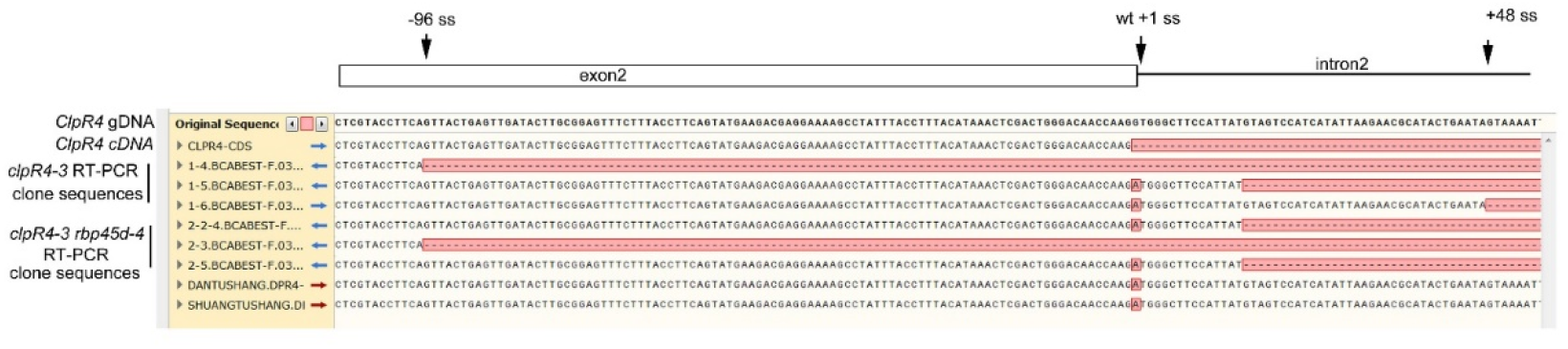
Sequences of the splicing products of *clpR4-3* and *clpR4-3 rbp45d-4.* The DNA sequences from different clones are aligned by SnapGene software. The sequences nearing *clpR4-3* intron 2 are showed.

**Figure S9.**
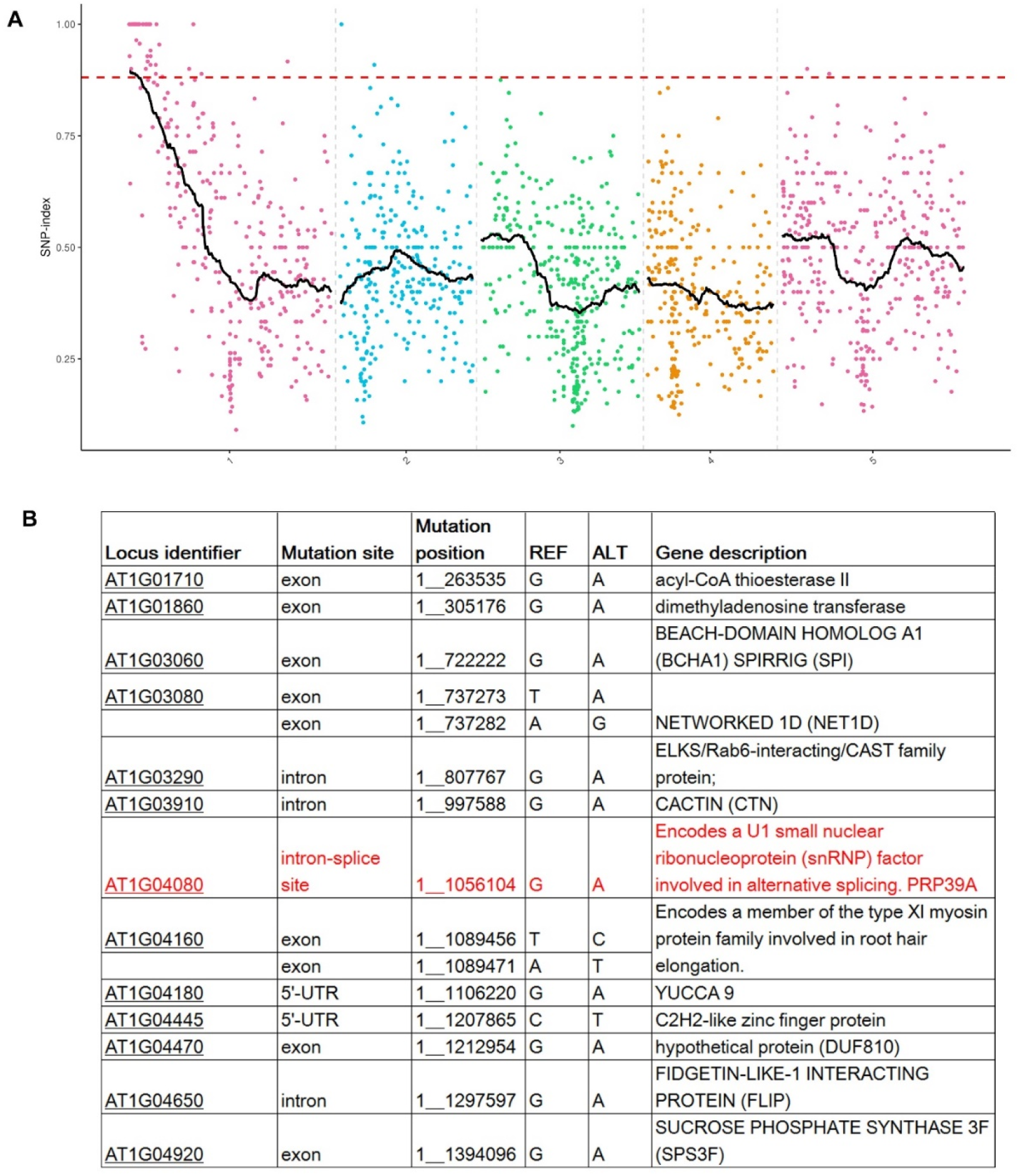
Cloning the *E25* suppressor gene through Mutmap technique. The BC_1_ F_2_ population was generated via a cross between *E25* and *sot5*. The DNA of thirty *E25* plants from the BC_1_ F_2_ population were extracted individually and pooled for Whole Genome Resequencing. (A) SNP index plots of five chromosomes of the *E25* mutant. Colored points indicate SNP positions and their indices. Black lines are regression lines. (B) *E25* gene was mapped in a region (~1.3 M) on Chromosome 1 which contains 15 candidate genes. Among them, at1g04080 which encodes PRP39a was the most likely candidate gene.

**Figure S10.**
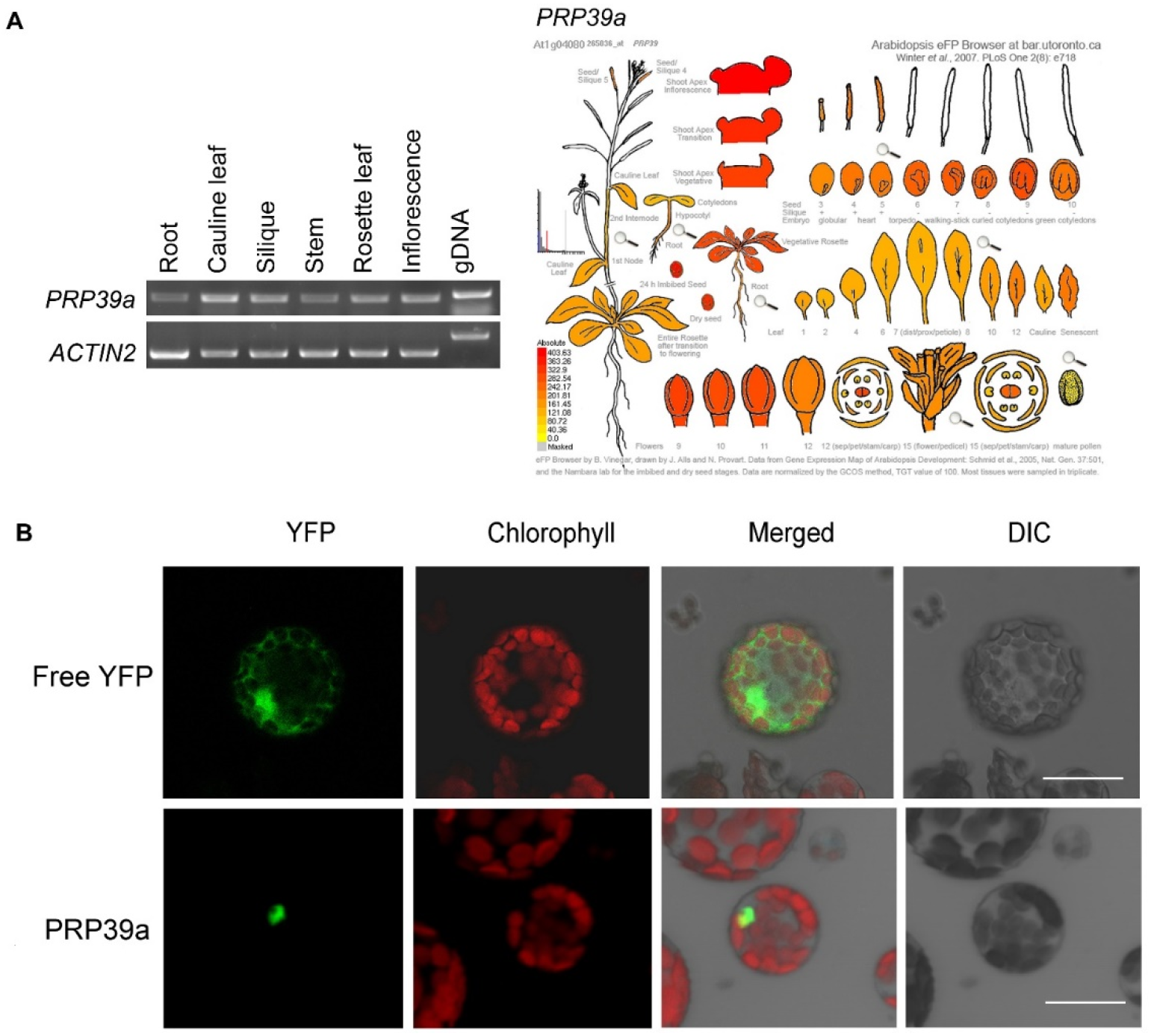
Expression pattern of *PRP39a* and subcellular localization of PRP39a protein. (A) RT-PCR analysis and eFP browser (http://bar.utoronto.ca/efp_arabidopsis/cgi-bin/efpWeb.cgi) show the expression pattern of *PRP39a* gene. (B) Subcellular localization of PRP39a. Microscopy analysis of the YFP fusion proteins transiently expressed in Arabidopsis protoplasts. The green fluorescence of the proteins was overlapped with chloroplast autofluorescence in merged images. DIC, Differential interference contrast microscopy. Bars = 15 μm.

**Figure S11.**
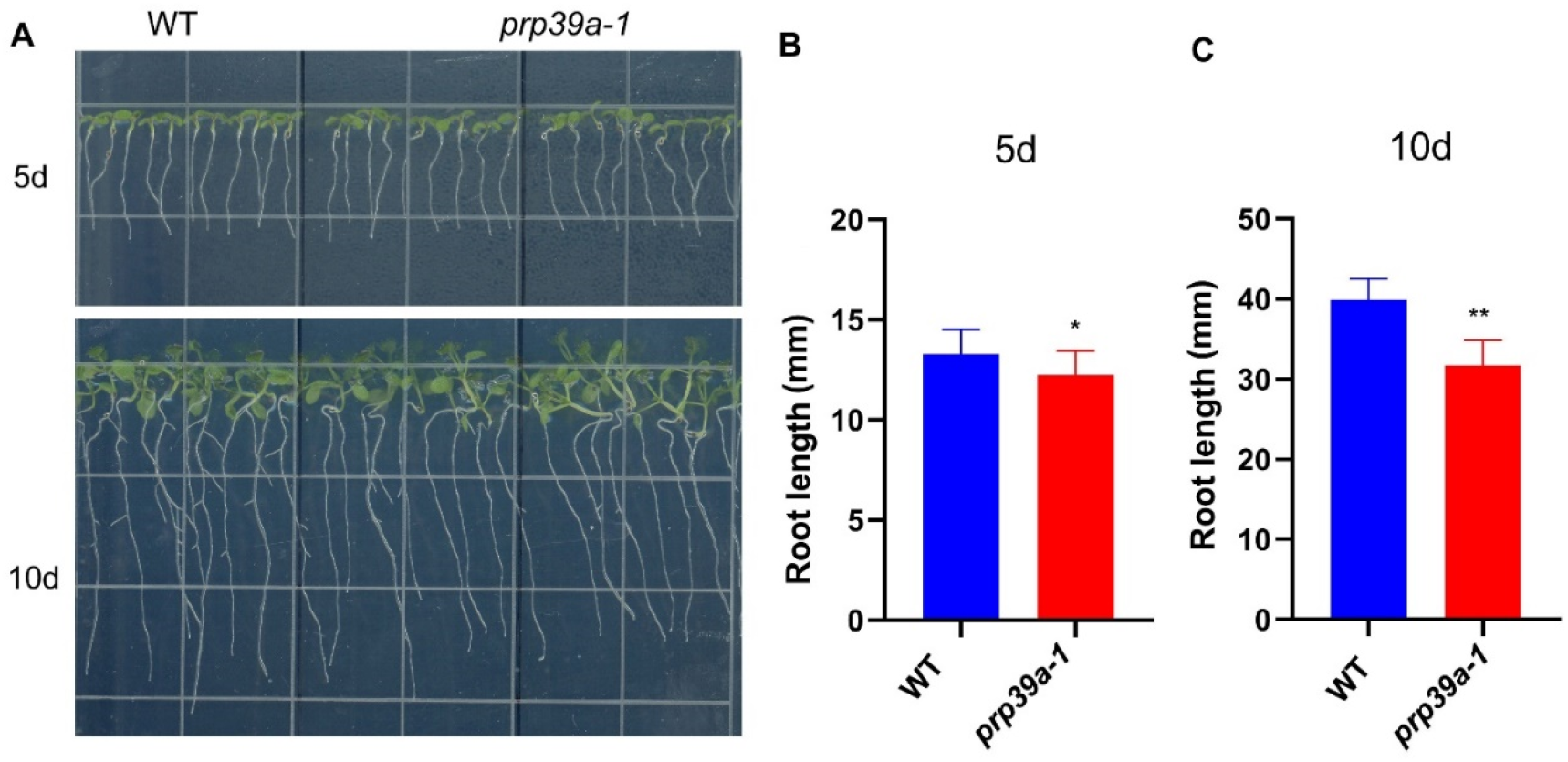
The *prp39a-1* mutant also exhibits the short primary root phenotype. (A) Short primary root phenotype of the 5d- and 10d-old *prp39a-1* seedlings. (B) and (C) are root length of 5d- and 10d-old seedlings. The data shown are means ± SD (n = 20). Student’s *t*-test; *, *P* < 0.05; Bar = SD. The experiments were performed at least two times.

